# Sequestration of the exocytic SNARE Psy1 into multiprotein nodes reinforces polarized morphogenesis in fission yeast

**DOI:** 10.1101/2020.05.01.072553

**Authors:** Kristi E. Miller, Joseph O. Magliozzi, Noelle A. Picard, James B. Moseley

**Affiliations:** Department of Biochemistry and Cell Biology, The Geisel School of Medicine at Dartmouth, Hanover, NH 03755

**Keywords:** Cell morphology, polarity, exocytosis, *Schizosaccharomyces pombe*, Psy1, Skb1

## Abstract

Polarized morphogenesis is achieved by targeting or inhibiting growth at distinct regions. Rod-shaped fission yeast cells grow exclusively at their ends by restricting exocytosis and secretion to these sites. This growth pattern implies the existence of mechanisms that prevent exocytosis and growth along non-growing cell sides. We previously identified a set of 50-100 megadalton-sized node structures along the sides of fission yeast cells that contain the interacting proteins Skb1 and Slf1. Here, we show that Skb1-Slf1 nodes contain the syntaxin-like SNARE Psy1, which mediates exocytosis in fission yeast. Psy1 localizes in a diffuse pattern at cell tips where it likely promotes exocytosis and growth, but Psy1 is sequestered in Skb1-Slf1 nodes at cell sides where growth does not occur. Mutations that prevent node assembly or inhibit Psy1 localization to nodes lead to aberrant exocytosis at cell sides and increased cell width. Genetic results indicate that this Psy1 node mechanism acts in parallel to actin cables and Cdc42 regulation. Our work suggests that sequestration of syntaxin-like Psy1 at non-growing regions of the cell cortex reinforces cell morphology by restricting exocytosis to proper sites of polarized growth.

## INTRODUCTION

Cell polarization is critical for the function of nearly every cell type and underlies essential processes such as cell growth and division. Regardless of how elaborate or simple a cell shape, polarized morphogenesis is achieved by targeting growth to specific regions. At the same time, polarized morphogenesis requires separate mechanisms that inhibit growth at other regions, thereby restricting growth to defined sites of polarity (Goehring and Grill, 2013). Fission yeast cells exhibit a highly polarized pattern of growth, making them an ideal model to study morphology and polarity. These rod-shaped cells maintain a constant cell width and grow exclusively from their tips by restricting exocytosis and secretion to these sites during interphase. During division, fission yeast redirect polarized exocytosis and secretion to the cell middle for septation and cell separation (Mitchison and Nurse, 1985; Kelly and Nurse, 2011; Das *et al*., 2007).

The growth machinery at cell tips has been widely studied (Reviewed by Martin and Arkowitz, 2013; Chiou *et al*., 2017). Landmark proteins such as Tea1 and Tea4 are deposited at cell ends by microtubules (Mata and Nurse, 1997; Feierbach *et al*., 2004; Martin *et al*., 2005; Tatebe *et al*., 2005). Landmark proteins recruit polarity factors including the formin For3 and its regulators, which assemble actin cables oriented toward cell tips (Feierbach *et al*., 2001; Martin *et al*., 2005; Martin and Chang, 2006; Martin *et al*., 2007). Actin cables act as tracks for myosin-based delivery of secretory vesicles, leading to targeted exocytosis of cell wall proteins and modifying enzymes at cell tips (Pruyne *et al*., 2004). At cell tips, a multiprotein complex called the exocyst tethers secretory vesicles for subsequent membrane fusion and content release mediated by SNAREs (soluble N-ethylmaleimide-sensitive factor-attachment protein receptors) (TerBush *et al*., 1996; Südhof and Rothman, 2009; Polgar and Fogelgren, 2018; Ganesan *et al*., 2020). SNARE proteins mediate membrane fusion in yeast and human cells, thus providing specificity by ensuring that only correctly targeted vesicles fuse (Protopopov *et al*., 1993; Sollner *et al*., 1993; Rothman, 1994; Pelham, 1999). Actin cables and the exocyst act in parallel to promote growth specifically at the tips of fission yeast (Bendezu and Martin, 2011; Snaith *et al*., 2011).

Mechanisms that promote growth at cell tips need to be reinforced by separate mechanisms that inhibit growth at cell sides. Previous studies identified Rga4 and Rga6, which are inhibitory GAPs (GTPase activating proteins) for the Rho GTPase Cdc42, along cell sides (Das *et al*., 2007; Tatebe *et al*., 2008; Kelly and Nurse, 2011; Revilla-Guarinos *et al*., 2016). These GAPs help to restrict Cdc42 activation and polarized growth to cell tips, but it has been unclear if other mechanisms exist to prevent exocytosis along cell sides. Here, we show that the fission yeast syntaxin-like SNARE protein Psy1 has different localization patterns at cell tips versus along cell sides. Psy1 forms a diffuse band at growing cell tips but localizes as cortical puncta along the non-growing cell sides. These puncta are the previously identified megadalton-sized node structures formed by the interacting proteins Skb1 and Slf1. We show that nodes sequester Psy1 at non-growing cortical sites to restrict exocytosis to proper sites of polarized growth.

## RESULTS

### Psy1 forms growth-positioned nodes

We noticed the presence of punctate structures in some images of Psy1 localization from previous studies (Wang *et al*., 2016; Zhu *et al*., 2018). Though typically considered a diffuse marker for the plasma membrane (PM), these images raised the possibility of a more intricate localization pattern. Therefore, we imaged GFP-Psy1 and collected series of z-sections to capture the full cell volume. At growing cell tips, Psy1 localized as diffuse bands at the PM. In contrast, along the non-growing cell middle Psy1 localized to discrete puncta on the PM (Figure 1A, left GFP-Psy1 panel). We observed these same Psy1 puncta using epifluorescence, laser-scanning confocal, and spinning disk confocal microscopy, so they do not represent an artifact of any particular imaging system. The distribution of Psy1 puncta suggested a link to cellular growth patterns. Psy1 puncta were excluded from sites of cell growth marked by the actin probe Lifeact-mCherry (Figure 1A). More specifically, Psy1 puncta were excluded from one end of small monopolar cells during interphase. In bipolar cells, Psy1 puncta were present at the cell middle and did not overlap with cortical actin patches at either growing end. In dividing cells, when actin and growth are redirected to the cell middle, Psy1 puncta were absent from the division septum. We conclude that Psy1 localizes in puncta at nongrowing regions of the cell cortex.

**Figure 1:**
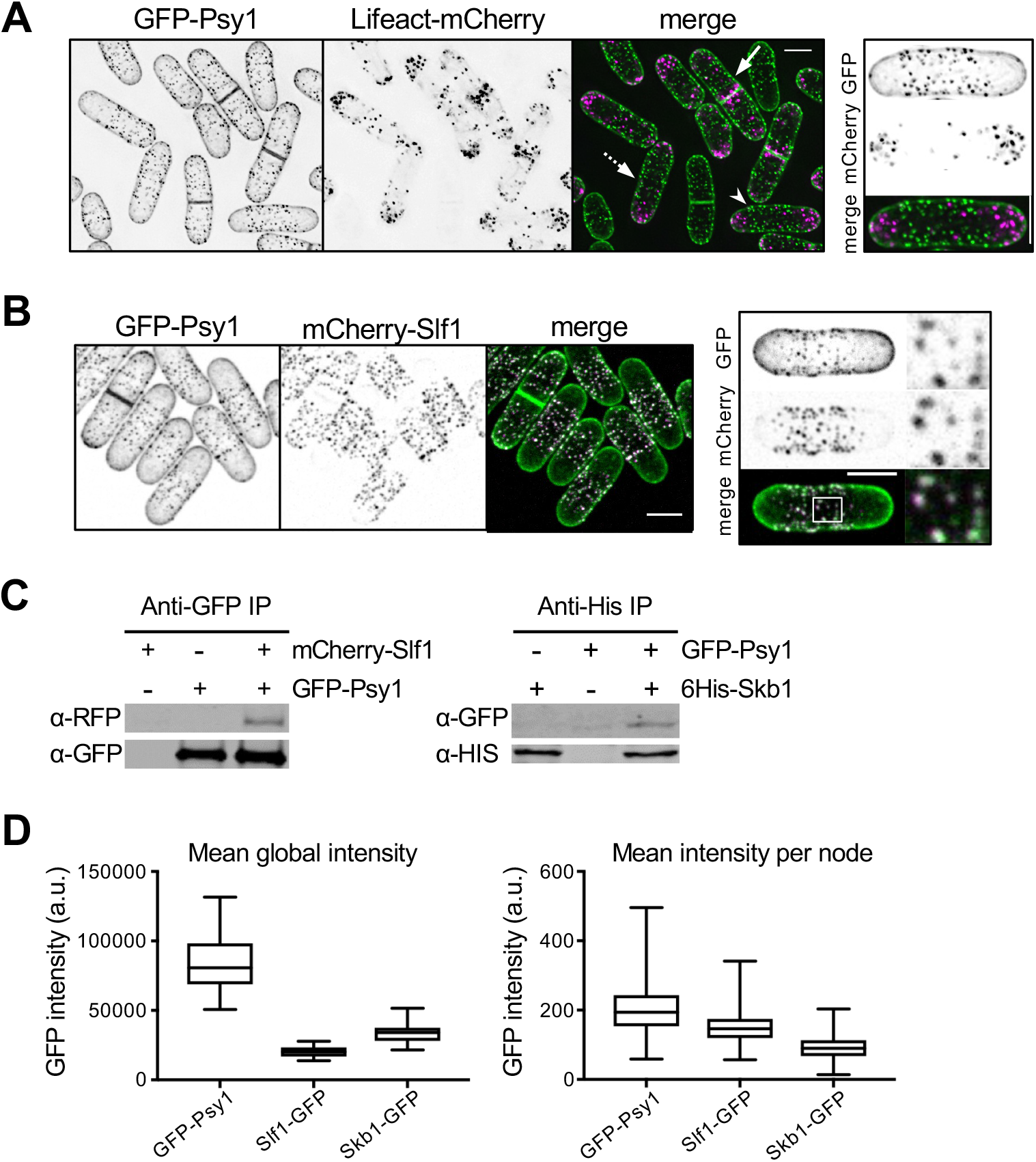
Psy1 localizes to Skb1-Slf1 nodes at non-growing regions of the PM. (A) GFP-Psy1 nodes do not overlap with actin marker Lifeact-mCherry. Images are from deconvolved z-series. Dashed arrow marks monopolar cell; arrowhead marks bipolar cell; arrow marks dividing cell. Representative image of bipolar cell is shown (Right) (B) Co-localization of GFP-Psy1 and mCherry-Slf1. Representative images with boxed region of close-up of mCherry and GFP node signals (Right) are shown. (C) Left, Co-immunoprecipitation of mCherry-Slf1 and GFP-Psy1 from fission yeast cells. Right, Co-immunoprecipitation of 6His-Skb1 and GFP-Psy1. (D) Global and local quantification (A.U.) of GFP-tagged Psy1, Skb1, and Slf1 proteins. Graph shows median as a line, quartiles, max, and min.

### Psy1 is a component of Skb1-Slf1 nodes

Psy1 puncta bear striking resemblance to a set of cortical node structures that we identified in previous studies (Deng and Moseley, 2013; Deng *et al*., 2014). These megadalton-sized nodes contain two interacting proteins called Skb1 and Slf1. We examined co-localization of Psy1 with these node proteins to test the possibility that Psy1 might be a component of Skb1-Slf1 nodes. In cells expressing mCherry-Slf1 and GFP-Psy1, we found that Psy1 and Slf1 co-localize in the same cortical nodes (Figure 1B). In addition to co-localization, Psy1 co-immunoprecipitated with both Skb1 and Slf1 (Figure 1C). Thus, Psy1 localizes to Skb1-Slf1 nodes and physically associates with these proteins. These results indicate that cortical nodes are multiprotein structures containing Skb1, Slf1, and Psy1.

We sought quantitative insight into the relationship between Skb1, Slf1, and Psy1 at nodes. By quantitative fluorescence microscopy, Skb1 and Slf1 are roughly stoichiometric at nodes, with each node containing on average 69 Skb1 molecules and 77 Slf1 molecules (Deng *et al*., 2014). The global concentration of GFP-Psy1 in cells was higher than either Skb1 or Slf1. At nodes, we measured slightly higher average signal for GFP-Psy1 than for Slf1 or Skb1 (Figure 1D). Based on these data, we estimate that each node contains ∼100 molecules of Psy1, although this number is likely to fluctuate between different nodes.

We tested if Psy1 localization to nodes requires Skb1 and/or Slf1. GFP-Psy1 localization to nodes was lost in either *skb1*Δ or *slf1*Δ mutant cells (Figure 2A). In both mutants, GFP-Psy1 localized in a diffuse pattern along the PM. This result is consistent with the interdependence of Skb1 and Slf1 for node formation (Deng *et al*., 2014). The absence of nodes led to an increased and largely constant concentration of diffuse Psy1 along cell sides, as well as increased levels of Psy1 at cell tips (Figures 2B-C). We conclude that Skb1 and Slf1 associate to form cortical nodes, which then recruit Psy1 to these structures through physical interactions. The presence of nodes reduces the concentration of diffuse Psy1 along cell sides.

**Figure 2:**
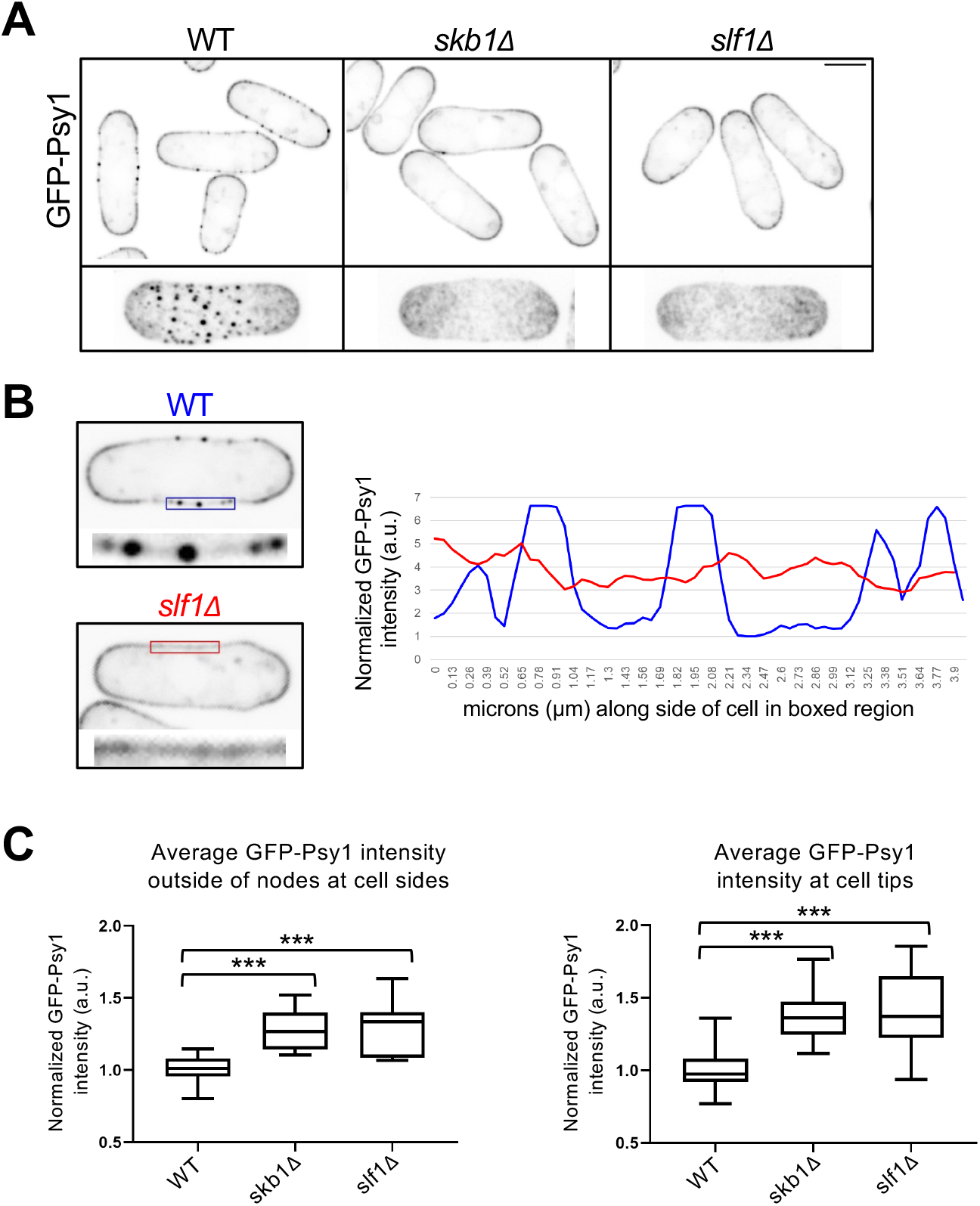
Skb1 and Slf1 are required for Psy1 localization to nodes. (A) (Top) Single middle z-slice images are shown. (Bottom) Maximum intensity projections of z-series are shown for a representative cell. Note the absence of Psy1 nodes in *skb1*Δ and *slf1*Δ cells. Scale bars 4µm. (B) Line scan of GFP-Psy1 intensity along the cell side of WT and *slf1*Δ cell (Right). Single middle z-slice images are shown with boxed region where line was drawn (Left). (C) Quantification of GFP-Psy1 intensity (A.U.) outside of nodes at cell sides (Left) and at cell tips (Right). n≥30 cells per strain. Graph shows median as a line, quartiles, max, and min.

### Psy1 protein is trapped in nodes

Next, we examined the dynamics of Psy1 at nodes. We performed fluorescence recovery after photobleaching (FRAP) of nodes along the cell side of WT cells. A bleached GFP-Psy1 region containing nodes did not recover fluorescence after 18 minutes (Figure 3A, left panel), similar to FRAP results from bleaching Psy1 along cell sides in previous studies (Bendezu *et al*., 2015; Wang *et al*., 2016; Tay *et al*., 2019). This result indicates that Psy1 protein is trapped in a node and does not exchange freely with Psy1 outside the node. In contrast, diffuse Psy1 along the cell side of *skb1*Δ and *slf1*Δ mutant cells was dynamic, similar to that of Psy1 at growing cell tips (Figures 3A-B). These findings were supported by time-lapse imaging of Psy1. In WT cells, we observed dynamic changes in Psy1 distribution at cell tips, but Psy1 at nodes did not move or rapidly disassemble over time (Figure 3C), similar to Skb1 and Slf1 (Deng and Moseley, 2013; Deng *et al*., 2014). However, Psy1 exhibited dynamic changes in its localization to cell tips and cell sides of *skb1*Δ and *slf1*Δ mutant cells.

**Figure 3:**
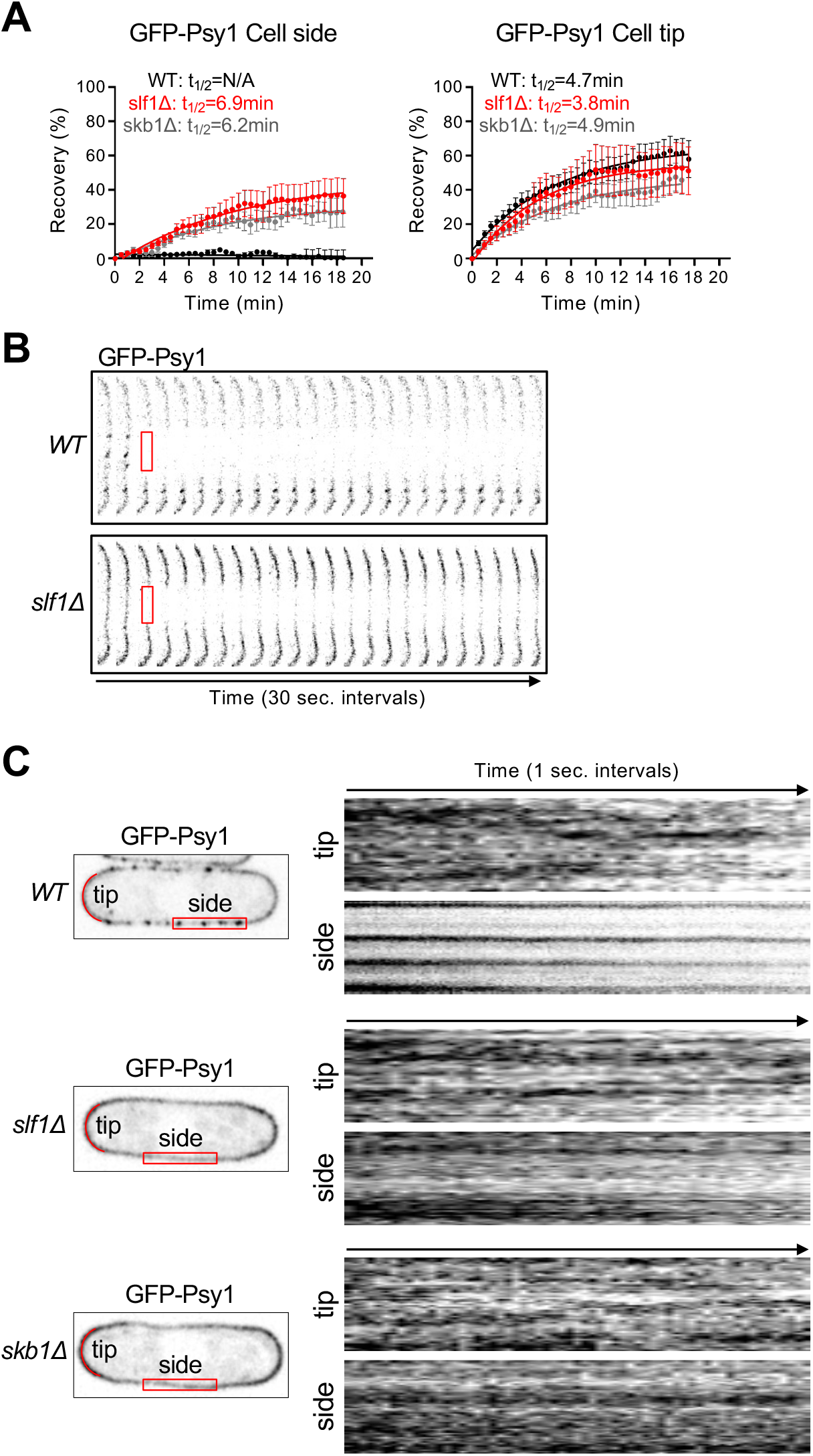
Psy1 nodes are stable structures. (A) FRAP curves of GFP-Psy1 recovery at the cell side (left) or at cell tips (right). n≥10 cells per strain. (B) Representative WT and *slf1*Δ cell sides before and after photobleaching. The red boxed region was bleached. (C) Time-lapse imaging of GFP-Psy1. Kymographs (Right) show distribution of GFP-Psy1 in red boxed region at the cell side, or along red lines at cell tips over a 70-second time period.

As an additional test of Skb1-Slf1-Psy1 node stability, we treated cells with 1,6-hexanediol, which disrupts weak hydrophobic interactions (Patel *et al*., 2007; Ribbeck and Gorlich, 2002). This chemical disrupts P-body granules in fission yeast cells (Kroschwald *et al*., 2015), which we confirmed in control experiments (Figure S1A). Treatment of cells expressing mCherry-Slf1 and GFP-Psy1 with 10% 1,6-hexanediol or DMSO control did not have a dramatic effect on Skb1-Slf1-Psy1 node stability (Figure S1B). This result indicates that Skb1-Slf1-Psy1 nodes are held together by interactive forces distinct from condensates such as P-bodies. Overall, our experiments reveal that nodes are static structures that trap Psy1 protein at cell sides.

### Genetic interactions implicate Skb1-Slf1 nodes in exocytosis

What is the function of Psy1 localization at cortical nodes? Psy1 is predicted to be necessary for exocytosis and growth at cell tips where it localizes independently of node proteins. Psy1 stably associates with Skb1-Slf1 nodes at non-growing regions along cell sides. We hypothesized that nodes might sequester Psy1 to inhibit exocytosis and growth along the cell middle. As a first test, we examined genetic interactions between node mutants and exocytosis mutants. In both *skb1*Δ and *slf1*Δ mutants, Psy1 is not trapped at nodes and instead localizes diffusively throughout the PM including along cell sides (Figures 2-3). We combined *skb1*Δ or *slf1*Δ with several mutations affecting exocyst function: the temperature-sensitive (ts) exocyst subunit mutants *sec8-1* and *sec3-2*, deletion of non-essential exocyst subunit Exo70, and deletion of the non-essential exocyst activator Rho3 (Wang, H. *et al*., 2003; Wang *et al*., 2002; Bendezu *et al*., 2012). Additionally, we combined *skb1*Δ or *slf1*Δ with *for3*Δ, which abolishes actin cables that direct polarized transport of exocytic secretory vesicles (Feierbach and Chang, 2001; Nakano *et al*., 2002). We discovered synthetic growth defects for *skb1*Δ*exo70*Δ and *skb1*Δ*for3*Δ mutants, while *skb1*Δ suppressed the growth defects of *sec8-1, sec3-2*, and to a minor extent *rho3*Δ mutants (Figure S2). Most double mutants with *slf1*Δ showed similar but less pronounced phenotypes, likely reflecting overlapping but non-identical functions for Skb1 and Slf1.

Mutations that affect exocytosis cause cell separation defects because exocytosis contributes to formation and remodeling of the division septum (Wang H *et al*., 2002; Bendezu *et al*., 2012; Jourdain *et al*., 2012; Wang N *et al*., 2016). Thus, we assayed the septation index in these double mutants and found increased septation index for *skb1*Δ*exo70*Δ double mutants compared to single mutants alone (Figures 4A-B). In contrast, we observed decreased septation index for *skb1*Δ*sec3-2* and *skb1*Δ*sec8-1* double mutants compared to *sec8-1* or *sec3-2* alone, similar to growth suppression (Figure S2). *slf1*Δ showed similar but less pronounced phenotypes at higher temperatures (Figures 4A-B). The synthetic defects observed with s*kb1*Δ*exo70*Δ (or s*lf11*Δ*exo70*Δ) and *skb1*Δ*for3*Δ mutants suggest that Skb1 and Slf1 may share a function with Exo70 and For3 in spatial control of exocytosis. Overall, these genetic interactions suggest that Skb1-Slf1-Psy1 nodes function in regulating exocytosis.

**Figure 4:**
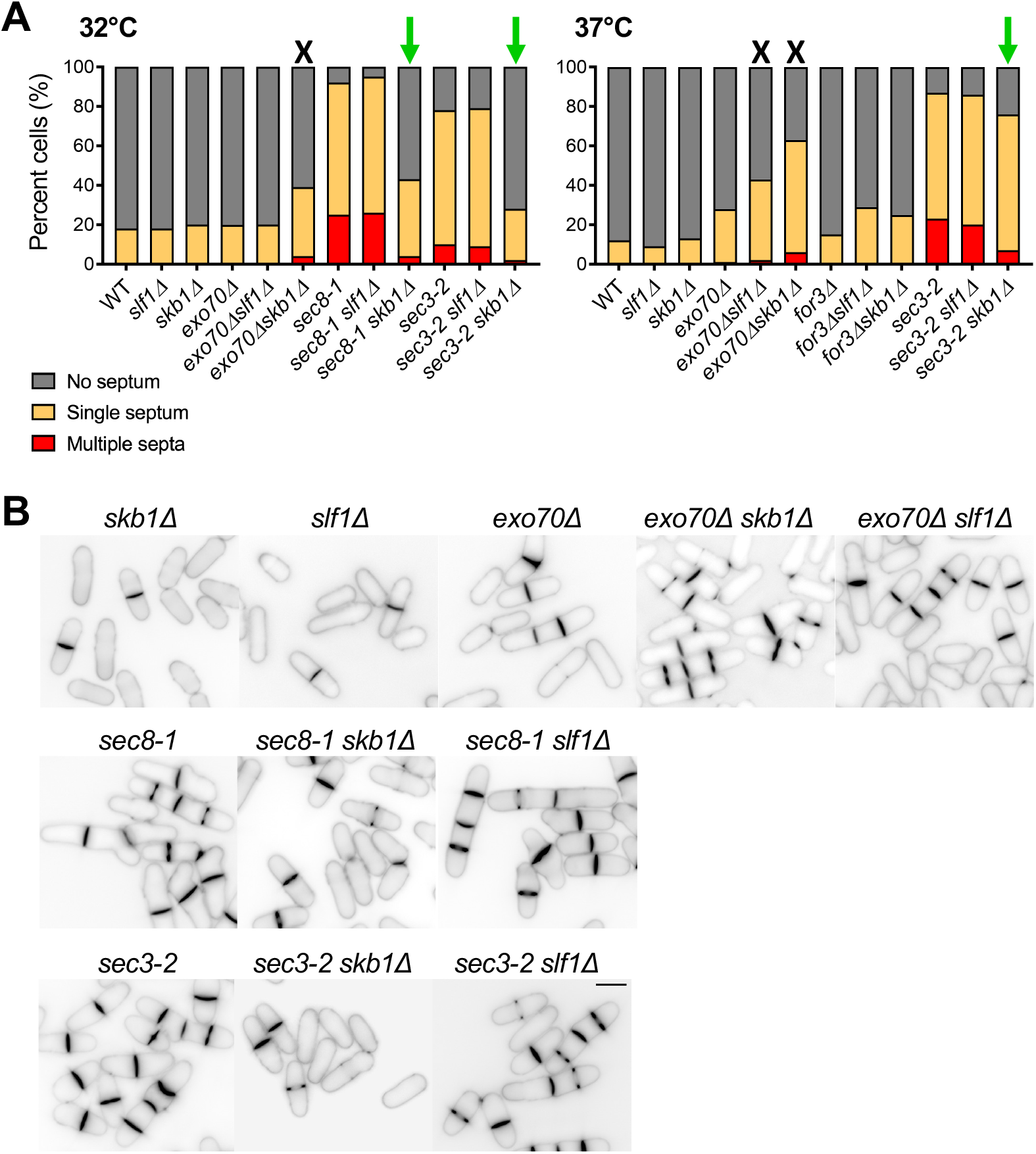
Genetic evidence for Skb1-Slf1-Psy1 node function in exocytosis. (A) Septation index in cells of indicated strains at 32°C (Left) or 37°C (Right). Growth defects or suppression with *skb1*Δ or *slf1*Δ is indicated by a black X or green arrow, respectively. n > 100 per strain. (B) Blankophor staining of indicated strains, showing cell separation defects of strains grown at 37°C. Bar, 7µm.

### Loss of Skb1-Slf1-Psy1 nodes leads to ectopic exocyst at cell sides

Based on these genetic interactions, we performed a series of microscopy-based experiments to test the role of Skb1-Slf1-Psy1 nodes in spatial control of exocytosis. Polarized exocytosis delivers the β-glucan synthase subunit Bgs4 to sites of active growth for cell wall synthesis (Cortes *et al*., 2005, 2015). In FRAP assays, photobleached GFP-Bgs4 signal at cell tips and the division site recovers rapidly (Figure 5A), consistent with cycles of endocytosis and polarized exocytosis at these sites. We found that *skb1*Δ and *slf1*Δ caused a significant decrease in the plateau of this recovery (Figure 5B), although the rate of recovery was unaffected. This result suggests that loss of nodes alters the normal trafficking and flux of Bgs4 protein at cell tips, consistent with Skb1-Slf1-Psy1 nodes contributing to spatial control of exocytosis in cells.

**Figure 5:**
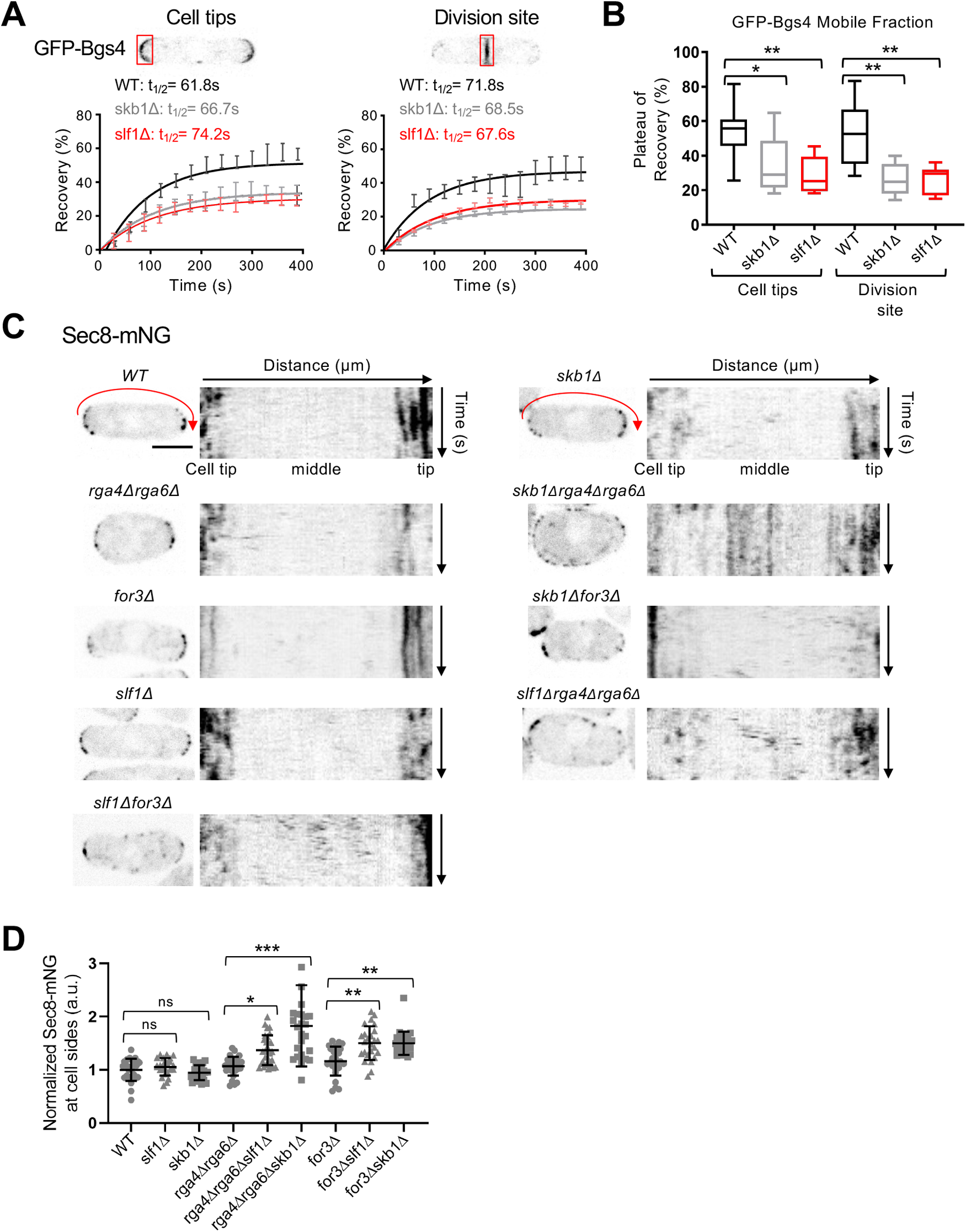
Loss of Skb1-Slf1-Psy1 nodes leads to changes in the pattern of exocytosis. (A) FRAP curves of GFP-Bgs4 at cell tips or the division site. Representative images of cells expressing GFP-Bgs4 are shown. n > 10 cells for each strain. (B) Plateau of recovery (%) of GFP-Bgs4 is reduced in *skb1*Δ and *slf1*Δ cells. *p = 0.01 and **p < 0.001 determined by ANOVA. Graph shows median as a line, quartiles, max, and min. (C) Localization of Sec8-mNG in the indicated strains. To the right of each single cell image is a kymograph showing Sec8 localization along cell sides (red arrow) over time. Scale bar, 4 µm. (D) Normalized mean intensity of Sec8-mNG along cell sides for the indicated strains. n > 25 for each strain. Graph shows mean ± SD. ns p ≥ 0.05, *p < 0.04, **p < 0.002, and ***p < 0.0001 determined by ANOVA.

To test this possibility more rigorously, we directly imaged exocyst component Sec8. Sec8-mNeonGreen (mNG) localized to growing cell tips and to the cell division site of WT cells as well as in *skb1*Δ and *slf1*Δ cells (Figure 5C). Because disruption of Skb1-Slf1-Psy1 nodes did not lead to aberrant exocyst localization, we tested the possibility that nodes act in parallel with other mechanisms to prevent exocytosis along cell sides. For example, the Cdc42 GAPs Rga4 and Rga6 localize along cell sides to prevent Cdc42 activation at these sites (Das *et al*., 2007; Tatebe *et al*., 2008; Kokkoris *et al*., 2014; Revilla-Guarinos *et al*., 2016). In time-lapse images of cells lacking both nodes and Cdc42 GAPs, we observed ectopic localization of the exocyst component Sec8-mNG in the middle of cells (Figures 5C). Importantly, these defects were more severe in *skb1*Δ *rga4*Δ *rga6*Δ (or *slf1*Δ *rga4*Δ *rga6*Δ) triple mutant cells than in *skb1*Δ (or *slf1*Δ) or *rga4*Δ *rga6*Δ mutants alone (Figure 5D). To extend this result, we performed similar assays for cells lacking nodes and actin cables, which target secretory vesicles away from cell sides. Combining node and actin cable mutations in the *skb1*Δ *for3*Δ (or *slf1*Δ *for3*Δ) double mutant led to aberrant localization of Sec8-mNG at cell sides (Figures 5C-D). Together, these results show that nodes sequester Psy1 to prevent mislocalization of the exocytic machinery to cell sides and act in parallel with other mechanisms including Cdc42 GAPs and actin cables.

### Skb1-Slf1-Psy1 nodes help maintain cell width

The spatial pattern of exocytosis defines fission yeast cell morphology, as localized secretion of cell wall proteins and modifying enzymes leads to cell growth (Bendezú and Martin, 2011; Atilgan *et al*., 2015; Abenza *et al*., 2015). Our results show that Skb1-Slf1-Psy1 nodes inhibit exocytosis at cell sides together with additional mechanisms mediated by Rga4, Rga6, and For3. Defects in spatial control of exocytosis should be accompanied by morphological consequences in cell shape. We found that cells lacking Skb1 were slightly wider than WT cells, but this difference was not statistically significant. Previous studies identified a cell width defect for *rga4*Δ, *rga6*Δ, and *for3*Δ mutants (Feierbach and Chang, 2001; Das *et al*., 2007; Revilla-Guarinos *et al*., 2016). Since *skb1*Δ and *slf1*Δ exhibited synthetic defects with these other mutants in positioning exocytosis, we tested their combined effects on cell morphology. We found that *skb1*Δ was additive with *rga4*Δ*rga6*Δ and *for3*Δ in increasing cell width, as the mutants with *skb1*Δ were significantly wider than any of the *rga4*Δ*rga6*Δ or *for3*Δ mutants alone (Figure 6). Cells lacking Slf1 showed similar results to *skb1*Δ but the effect was less pronounced (Figure 6), which may be due to additional functions for Skb1 as suggested by previously noted differences in *skb1*Δ and *slf1*Δ phenotypes (Deng *et al*., 2014). These combined results indicate that Skb1-Slf1 nodes sequester Psy1 at the cell middle to prevent aberrant exocytosis and this mechanism works in parallel to previously identified mechanisms including Cdc42 GAPs and actin cables. Consistent with Skb1-Slf1-Psy1 nodes functioning independently of Cdc42 GAPs at cell sides, nodes do not co-localize with Rga4 or Rga6 (Figure S3). Overall, these results suggest that Skb1-Slf1 nodes work in parallel with Cdc42 GAPs and actin cables for polarized morphogenesis.

**Figure 6:**
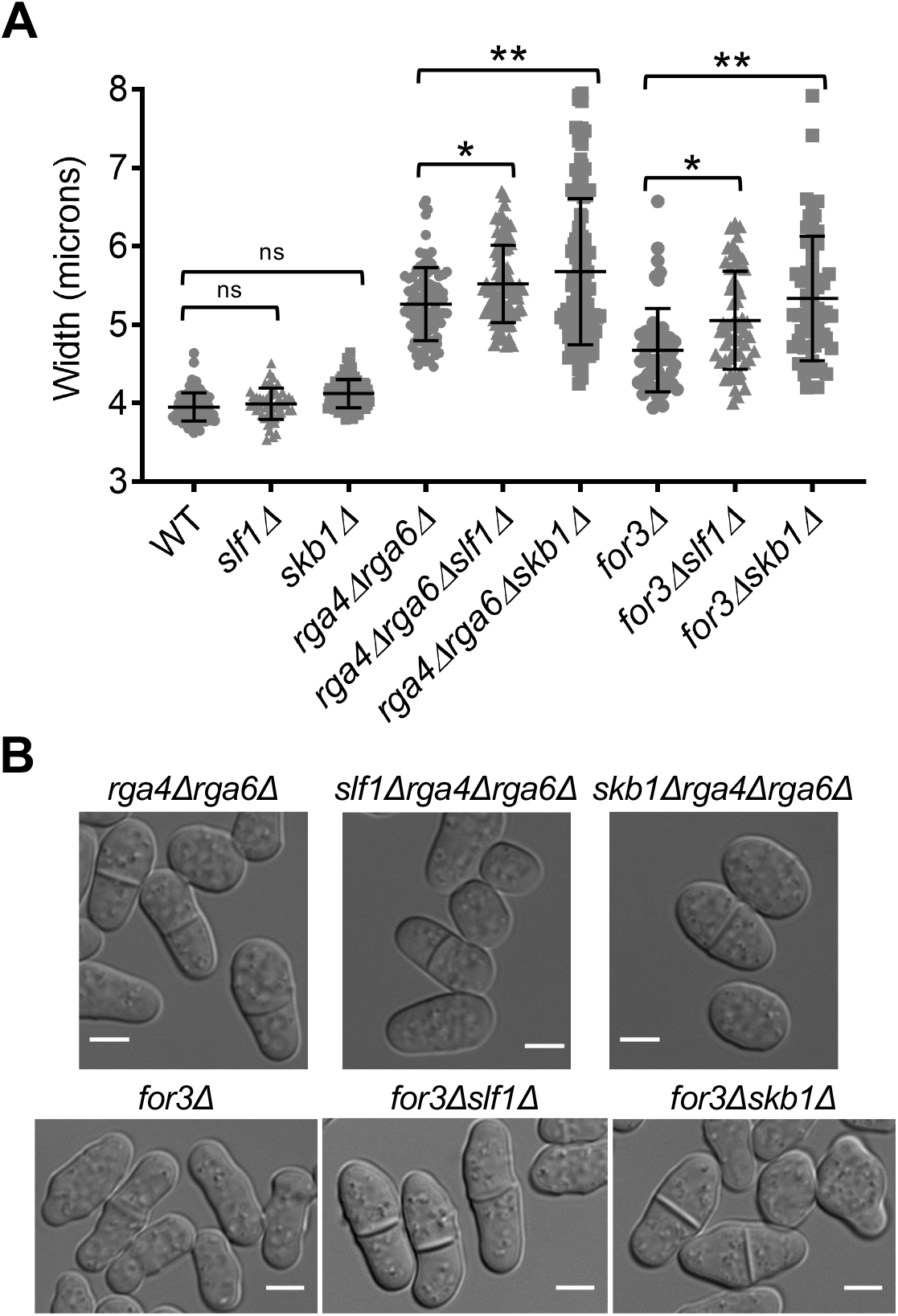
Nodes contribute to cell width control. (A) Cell width (µm) of indicated cell types. ns p ≥ 0.05, *p < 0.05, and **p < 0.0001 determined by ANOVA. n > 50 for each cell type. Graph shows mean ± SD. (B) Representative images showing wider cells with *skb1*Δ and *slf1*Δ. Bars, 4µm.

### Generation of a psy1 mutant that does not localize to Skb1-Slf1 nodes

Defects in exocyst localization and cell morphology in *skb1*Δ and *slf1*Δ mutants may be caused by loss of Psy1 sequestration at nodes, but we cannot exclude a role for other aspects of Skb1 and Slf1 function in these defects. To focus more specifically on Psy1 localization at nodes, we sought to generate a Psy1 mutant that no longer localizes to Skb1-Slf1 nodes. Because *psy1* is an essential gene, we initially examined truncated Psy1 constructs fused to GFP and integrated into the genome of cells that also expressed WT *psy1*. Truncated GFP-Psy1 lacking its N-terminal H_abc_ domain (H_abc_Δ) or C-terminal membrane anchor (MAΔ) no longer localized to nodes, while a Psy1 protein lacking its SNARE motif (SNAREΔ) retained node localization (Figure S4). psy1-MAΔ had an overall cytoplasmic distribution, consistent with a role in PM localization. Since psy1-H_abc_Δ retained PM localization while disrupting node localization, we focused on the role of the H_abc_ domain in node localization.

H_abc_ domains of SNARE proteins consist of three alpha-helical regions termed Ha, Hb, and Hc. We mutated charged surface-exposed residues found outside and within the alpha-helical regions of the H_abc_ domain (Figure 7A). Two mutants were identified with reduced localization to nodes: psy1-Ha-m1, which contains 5 mutations of charged residues to alanine in the N-terminal half of the Ha region; and psy1-Hc-m1, which contains 5 mutations of charged residues to alanine in the N-terminal half of the Hc region (Figures 7A-B; See Table S2 for specific amino acid mutations). psy1-Ha-m1 localized homogenously throughout the plasma membrane with minimal concentration at nodes. psy1-Hc-m1 localized homogenously throughout the plasma membrane but also ectopically to intracellular puncta, which were seen most clearly in middle focal plane microscopy images (Figure S5A). We chose the psy1-Ha-m1 mutant to characterize further because it lost node localization without additional changes to localization.

**Figure 7:**
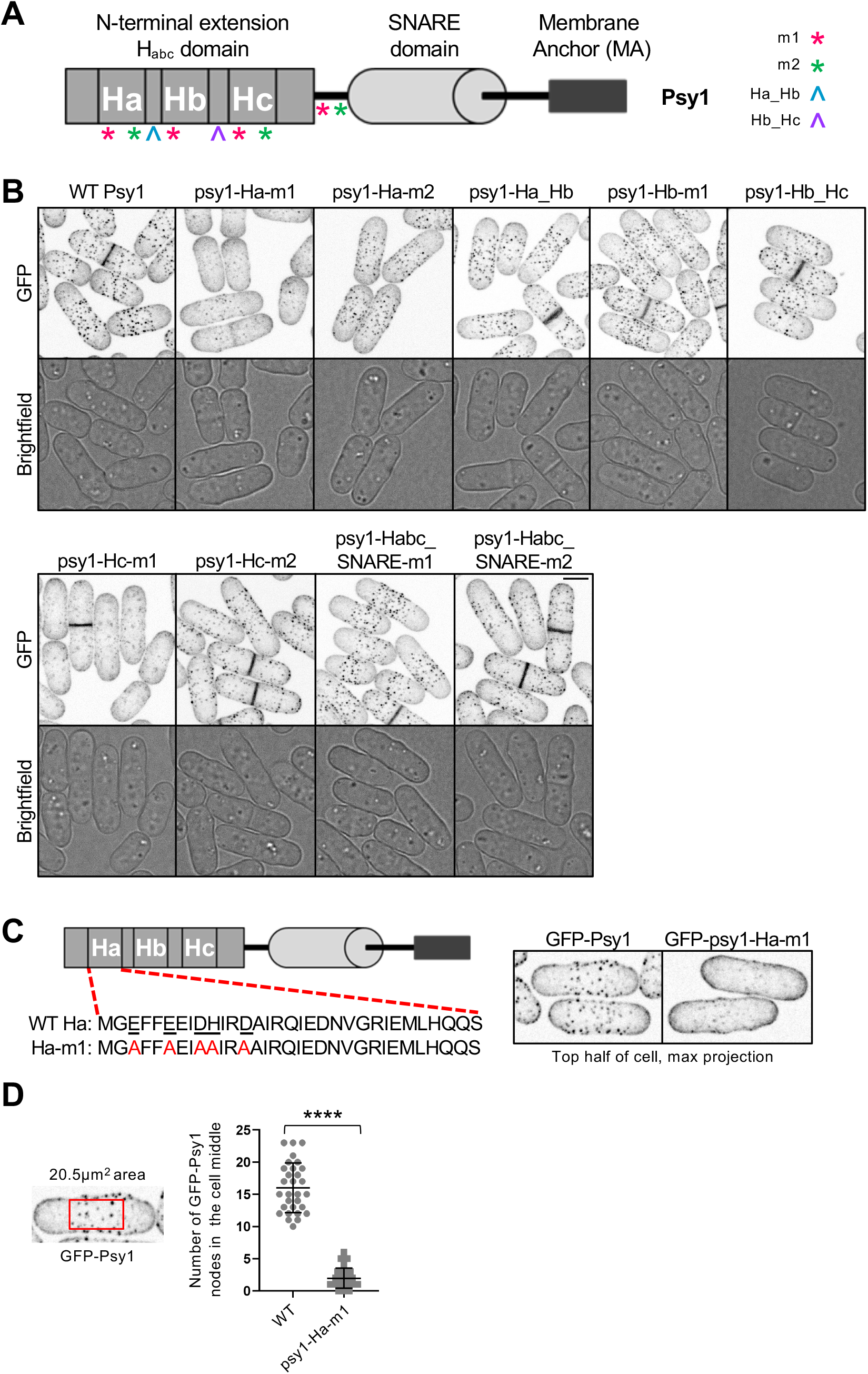
Analysis of Psy1 mutant node localization. (A) Diagram of Psy1 protein domains and notation of location of mutations (See Table S2 for specific amino acid mutations). (B) Maximum projection images of z-series taken of GFP tagged Psy1 or mutant form of psy1 as listed. Single middle z-slice images shown for brightfield. Bar, 4µm. (C) Amino acid sequence of Ha domain in WT Psy1 or psy1-Ha-m1 (Left). Representative images showing GFP tagged Psy1 or psy1-Ha-m1. Maximum intensity projections were created from the top half of cell z-stack images that encompass one side of the cell membrane (Right). (D) Number of GFP-Psy1 nodes in the cell middle above a set threshold intensity and size. **** p < 0.0001 determined by ANOVA. n≥30 for each cell type. Graph shows mean ± SD. (Left) Representative image showing size of 20.5µm^2^ boxed region that was used to define the middle of cells for analysis. The cell is reproduced from panel C to show selection of region of interest.

### Psy1 sequestration into Skb1-Slf1 nodes promotes polarized morphogenesis

From confocal images, GFP-psy1-Ha-m1 exhibited dramatically reduced node localization (Figure 7C). We quantified the number of Psy1 nodes in the cell middle using a threshold method and found that psy1-Ha-m1 localization to nodes was indeed significantly reduced compared to WT cells (Figure 7D). The global intensity of GFP-psy1-Ha-m1 was similar to wild type GFP-Psy1 (Figure S5B), meaning that loss of node localization was not due to changes in overall protein concentration. We conclude that the H_abc_ domain is required for Psy1 localization to nodes, and the psy1-Ha-m1 mutant fails to localize to nodes.

Using this new mutant, we tested whether Psy1 is necessary for Skb1-Slf1 node formation. The appearance, number, and intensity of mCherry-Slf1 nodes in the *psy1-Ha-m1* mutant was unchanged compared to WT cells (Figure S6), showing that Skb1-Slf1 nodes remain intact when Psy1 is not concentrated in these structures. This result supports a model where Skb1 and Slf1 form the required and interdependent core of nodes, and Psy1 is recruited as a peripheral component of these structures.

Next, we used the psy1-Ha-m1 mutant to test the functional role of Psy1 at nodes. For these experiments, we imaged exocyst component Sec8-tdTomato similarly to our earlier experiments with *skb1*Δ and *slf1*Δ. From time-lapse images of cells containing *psy1-Ha-m1* and lacking Cdc42 GAPs, we observed ectopic localization of Sec8-tdTomato in the cell middle (Figure 8A). Critically, this defect was greater in the *psy1-Ha-m1 rga4*Δ *rga6*Δ triple mutant compared to either *rga4*Δ *rga6*Δ double mutant or the *psy1-Ha-m1* mutant alone (Figure 8B). These results indicate that sequestration of Psy1 into nodes prevents mislocalization of exocytic machinery at cell sides.

**Figure 8:**
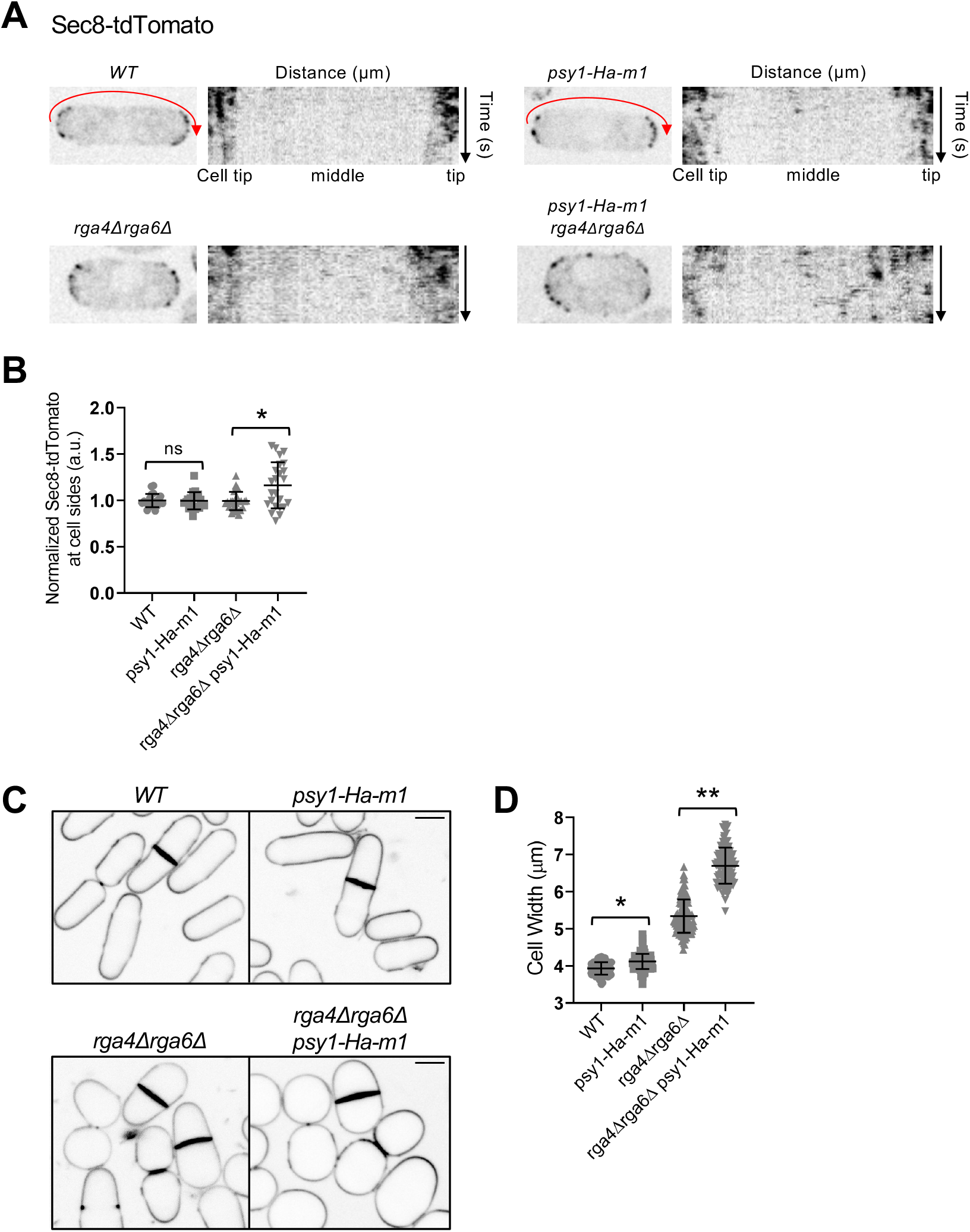
*psy1-Ha-m1* exhibits ectopic exocyst at cell sides and increased cell width. (A) Localization of Sec8-tdTomato in indicated strains. To the right of each single cell image is a kymograph showing Sec8 localization along cell sides (red arrow) over time. (B) Normalized mean intensity of Sec8-tdTomato along one cell side for the indicated strains. n > 25 for each strain. Graph shows mean ± SD. ns p ≥ 0.05 and *p < 0.001 determined by ANOVA. (C) Blankophor staining of indicated strains, showing increased cell width with *psy1-Ha-m1*. Bars, 4µm. (D) Cell width (µm) of indicated cell types. *p < 0.001 and ** p < 0.0001 determined by ANOVA. n > 100 for each cell type. Graph shows mean ± SD.

Defects in spatial control of exocytosis of mutants lacking Cdc42 GAPs and *skb1*Δ (or *slf1*Δ) were accompanied by morphological changes in cell shape. We found that *psy1-Ha-m1* cells were significantly wider than WT cells. Further, combining *psy1-Ha-m1* with *rga4*Δ *rga6*Δ dramatically increased cell width compared to either *rga4*Δ *rga6*Δ double mutant or the *psy1-Ha-m1* mutant alone (Figures 8C-D). We conclude that sequestration of Psy1 into Skb1-Slf1 nodes prevents aberrant exocytosis at cell sides to reinforce polarized morphogenesis in fission yeast.

## DISCUSSION

In this study, we have shown that Psy1 is sequestered in Skb1-Slf1 nodes at non-growing regions of the PM to prevent ectopic exocytosis at these sites. Our genetic experiments reveal that this new mechanism acts in parallel to actin cable transport and Cdc42 GAPs. This demonstrates that ectopic growth along cell sides is inhibited by multiple mechanisms that reinforce cell morphology, in particular for control of cell width (Figure 9). We anticipate that signaling pathways that control Psy1 nodes, actin cables, and Cdc42 GAPs are further reinforced by physical barriers to exocytosis at cell sides such as the cortical endoplasmic reticulum (Ng *et al*., 2018). This combination of mechanisms along cell sides supports the notion that cell polarity arises from multiple activities that affect different steps in the trafficking, docking, and fusion of exocytic secretory vesicles.

**Figure 9:**
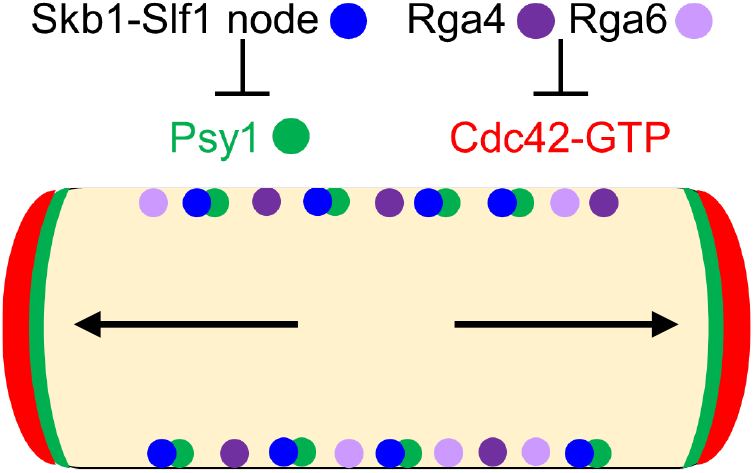
Working model for polarized morphogenesis through inhibition of exocytosis along the sides of fission yeast cells. Green circles denote Psy1 sequestered by Skb1-Slf1 nodes (blue circles) along cell sides. Rga4 and Rga6 (Purple circles) inhibit Cdc42-GTP at these sites. Cdc42-GTP denoted as a broad red band directs actin cable (black arrows) organization to cell tips. Cables act as tracks for secretory vesicles that fuse to the PM through the action of SNARE Psy1 (broad green band at tips) to facilitate growth.

We examined the functional role of Psy1 sequestration into nodes using three different mutants: *skb1*Δ, *slf1*Δ, and *psy1-Ha-m1*. Each of these mutants prevents Psy1 localization to nodes, providing three separate test cases for the function of Psy1 at nodes. In all three cases, we observed ectopic exocytosis along cell sides and an increase in cell width, particularly when combined with mutations in Cdc42 GAPs. The combination of results from these three mutants strongly supports the model that nodes sequester Psy1 to prevent exocytosis along cell sides. We note that the severity of phenotypes varies among these three mutants, likely reflecting differences in the functions and protein-protein interactions for Skb1 and Slf1. In addition, we do not exclude the possibility that the *psy1-Ha-m1* mutation alters the activity of Psy1. In fact, some members of the Syntaxin family undergo intramolecular conformational changes that regulate their activity and vesicle fusion, where the H_abc_ domain folds back onto the SNARE motif creating a ‘closed’ and inactive conformation (Dulubova *et al*., 1999; Gerber *et al*., 2008; MacDonald *et al*., 2010). More work is needed to test if Psy1 exhibits such an autoinhibitory conformation, as well as the role of the Ha-m1 mutation in this mechanism. More generally, it will be interesting to learn if nodes regulate Psy1 activity in addition to its localization through interactions with Skb1, Slf1, or possibly via other unidentified node proteins.

Sequestration of Psy1 into nodes by Skb1-Slf1 has implications for SNARE proteins beyond yeast. We note that SNARE proteins have been shown to be clustered and sequestered in human cells with connection to several diseases. For example, individuals with insulin resistance exhibit sequestration of SNAP-23 (synaptosomal-associated protein of 23 kDa) by lipid droplets, thus blocking exocytosis of glucose transporters (Boström *et al*., 2007). In addition, SNAREs are aberrantly sequestered in cholesterol-enriched regions of LSD (lysosomal storage disorder) endolysosomal membranes. This abnormal spatial organization locks SNAREs in complexes and impairs their sorting and recycling (Fraldi *et al*., 2010). Alpha-synuclein aggregates preferentially sequester SNAP-25 and VAMP-2 (vesicle-associated membrane protein 2) leading to reduced exocytosis and contribute to neurotoxicity (Choi *et al*., 2018). Overall, regulated sequestration of SNARE proteins may be a general mechanism leading to inhibition of exocytosis in a wide range of cell types and organisms.

## MATERIALS AND METHODS

### Strains, plasmids, and growth conditions

Standard methods were used to grow and culture *S. pombe* cells (Moreno *et al*., 1991). Yeast strains used in this study are listed in Supplemental Table S1. Gene fusions were expressed from their endogenous promoters unless otherwise indicated. One-step PCR-based homologous recombination was performed for tagging or deletion of genes on the chromosome (Bähler *et al*., 1998). To obtain N-terminal GFP-tagged Psy1, the *psy1+* promoter was amplified using primers containing *Bgl*II and *Pac*I sites at the 5’ and 3’ends, respectively. This PCR product was inserted in place of the *P3nmt1* promoter in the *pFA6A-kanMX6-P3nmt1-GFP* plasmid. PCR product from the *pFA6A-kanMX6-psy1+-GFP* plasmid was inserted adjacent (at the 5’ end) to the *psy1* open reading frame on the chromosome. Slf1 N-terminal mCherry tagging was described previously (Deng *et al*., 2014). Plasmids used in this study are listed in Supplemental Table S2. The majority of plasmids used in this study contain the pDC99 backbone for integration at the *leu1* locus and include the *ura4+* cassette. To create pDC99-Ppsy1-GFP-psy1-Tpsy1 we PCR amplified Ppsy1-GFP-psy1-Tpsy1 using 5’ and 3’ primers containing *Kpn*1 and *Sac*II sites, respectively, from genomic DNA of fission yeast cells. This PCR product was then inserted into *Kpn*1/*Sac*II digested pDC99 plasmid. Mutations or substitutions were introduced into *psy1* in the pDC99-Ppsy1-GFP-psy1-Tpsy1 plasmid using the NEB Q5 site directed mutagenesis kit.

To measure cell width at division (Figures 6 and 8), prototrophic strains were grown in EMM4S medium at 32°C and stained with Blankophor prior to imaging. To determine the septation index of cells (Figure 4A), cells were grown at 25°C in EMM4S medium and then cultures were shifted to 32°C or 37°C for 5h before staining with Blankophor and imaging. In Figure S1, cells were grown at 25°C in EMM4S medium and then treated with 10% 1,6-hexanediol or DMSO for 45 min prior to imaging.

### Co-immunoprecipitation and immunoblotting

Co-immunoprecipitation of proteins from fission yeast cell extracts were performed using a protocol adapted from Deng and Moseley, 2013. For Figure 1C, mCherry-Slf1 and GFP-Psy1 co-immunoprecipitation was performed using 50 mL of 0.5 OD_595_ cells grown in YE4S at 25°C.

Cells were harvested by centrifugation, washed 1x in Milli-Q water and resuspended in 400 μl of Lysis Buffer (20 mM HEPES (4-(2-hydroxyethyl)-1-piperazineethanesulfonic acid) pH 7.4, 200 mM NaCl, 1% Triton X-100, 1mM phenylmethylsulfonyl fluoride, complete EDTA-free protease inhibitor tablets [Roche, Indianapolis, IN]) together with 200 μl of glass beads and lysed using a Mini-beadbeater-16 (BioSpec, Bartlesville, OK; two cycles of 1 minute at max speed). Lysates were spun at 14,000 x *g* for 10 minutes at 4°C and supernatants were recovered. α-GFP magnetic beads (Allele Biotech) were washed 3x in Lysis Buffer and added to clarified cell lysates and rotated for 2 hours at 4°C. After lysate removal, beads were washed 5x in Lysis Buffer and resuspended in SDS-PAGE sample buffer (65 mM Tris, pH 6.8, 3% SDS, 10% glycerol, 10% 2-mercaptoethanol) and boiled for 5 minutes at 99°C followed by SDS-PAGE and western blotting.

For detection of 6His-Skb1 and GFP-Psy1 interaction, pJM482 (pREP3x-6His-skb1) or pJM210 (pRep3x) was transformed into appropriate strains and colonies were selected on EMM-Leu + 10 μg/mL thiamine plates. Cells were grown in 50 mL EMM-Leu + 10μg/mL thiamine until reaching a 0.5 OD_595_. Co-immunoprecipitation and SDS-PAGE followed by western blotting was carried out as mentioned above except TALON metal affinity resin (Takara) was used instead of α-GFP beads for 6His-Skb1 enrichment.

Western blots were probed with anti-6His (SC-8036; Santa Cruz Biotechnology), anti-GFP (Moseley et al., 2009), and anti-RFP (NBP1-69962; Novus Biologicals) antibodies.

### Microscopy and image analysis

Fission yeast cells were grown in EMM4S medium to logarithmic phase for imaging. Time-lapse imaging was performed using two spinning disk confocal microscope systems. The first system was an Andor CSU-WI (Nikon software) equipped with a 100x 1.45 NA CFI Plan Apochromat Lambda objective lens (Nikon); 405 nm, 445 nm, 488 nm, 560 nm, 637 nm and 685 nm laser lines; and a Andor Zyla camera on an inverted microscope (Ti-E, Nikon). The second system was a Yokogawa CSU-WI (Nikon Software) equipped with a 100x 1.45 NA CFI Plan Apochromat Lambda objective lens (Nikon); 405 nm, 488 nm, 561 nm laser lines; and a photometrics Prime BSI camera on an inverted microscope (Eclipse Ti2, Nikon). Experimental and control samples in a data set were always collected on the same microscope to ensure the accurate comparison of images for analysis.

For most time-lapse imaging, single z-slice images were captured every 1 or 3 sec using cells mounted on an agarose slab at room temperature. Static images of fission yeast cells shown in Figure 1A, 4B, 6B, S1, and S3 were taken at room temperature with a DeltaVision Imaging System (Applied Precision), equipped with an Olympus IX-71 inverted wide-field microscope, a Photometrics CoolSNAP HQ2 camera, and Insight solid-state illumination. Most images were acquired with 11 z-stacks and 0.4 μm step using cells mounted in EMM4S liquid media. Images were iteratively deconvolved using SoftWoRx software (Applied Precision), as indicated in figure legends. All other static images shown were taken at room temperature using the spinning disk confocal microscopy systems described above.

Image processing and analysis were performed using ImageJ (National Institutes of Health). Figures were generated using maximum intensity projections of z-stacks for fluorescent images and a single middle z-section for DIC images, unless otherwise indicated in the figure legend. For global intensity quantification of GFP tagged Psy1, Skb1, or Slf1 (Figure 1D), sum intensity projections were created from images (40 z-sections, 0.17 μm spacing), and an ROI was drawn around the outline of cells. The intensity of an equal sized ROI with no cell present was used to subtract background. For the mean intensity (integrated density) per node measurements (Figure 1D), sum projections were created from the top half of cell z-stack that encompass one side of the cell membrane. An ROI was drawn around each individual node (measured 175 nodes per strain) and background was subtracted using the intensity of an equal sized ROI with no node present. Images used to obtain these measurements were taken with a Zeiss LSM 880 laser scanning confocal microscope (See below).

To quantify mean Sec8-mNG (or Sec8-tdTomato) intensity in the cell middle over time (Figures 5D and 8B), the intensity of an equal sized ROI with no cell present was used to subtract background from selected cells. Using the segmented line tool, a line was drawn from the middle of cell tips along the side of the cells. Kymographs were generated using the multiple kymograph plug-in for ImageJ to display Sec8-mNG (or Sec8-tdTomato) signal over time along the line. The mean Sec8-mNG (or Sec8-tdTomato) intensity over time in the cell middle was measured from the kymograph. The same sized box was drawn to cover the same width along the cell side for all cell types. Intensity values were normalized to the average Sec8 intensity in the cell middle of WT cells and plotted on a graph.

To examine GFP-Psy1 intensity along the cell sides, the intensity along an equal sized line was analyzed from middle Z-plane images after background subtraction. Peaks of GFP-Psy1 intensity indicating the presence of a Psy1 node along the sides of WT cells were excluded from data points to obtain the average GFP-Psy1 intensity outside of nodes at cell sides. The average GFP-Psy1 intensity along cell sides for each cell was normalized to the average GFP-Psy1 intensity outside of nodes in WT cells and plotted on a graph (Figure 2C). Average GFP-Psy1 intensity at cell tips was determined from middle Z-plane images after background subtraction. A line was drawn at cell tips and the average GFP-Psy1 was determined. Values for each cell were normalized to the average GFP-Psy1 intensity at cell tips of WT cells and plotted on a graph (Figure 2C).

To quantify the number of GFP-Psy1 nodes in the cell middle (Figure 7D), maximum intensity projections of the top half of cell z-stack images (1-10z) that encompass one side of the cell membrane were analyzed after background subtraction. A 20.5µm^2^ rectangular region was used to define the middle of cells for analysis. A fluorescence threshold was set above background that selected fluorescent pixels at nodes. Next, the values above a threshold were analyzed by size and spots larger that 0.02µm^2^ were counted as nodes and plotted on a graph (Figure 7D).

For GFP-Psy1 global intensity quantification, sum intensity projections were created from all 17 z-sections at 0.3 μm spacing, and an ROI was drawn around the outline of cells. The intensity of WT cells without any fluorescently tagged proteins were used to subtract background. Values were normalized to the average GFP-Psy1 global intensity of WT cells and plotted on a graph (Figure S5B)

To quantify the number and intensity of mCherry-Slf1 nodes per cell, maximum intensity projections were created from all 17 z-sections at 0.3 μm spacing. Brightfield images were used to create a binary cell mask to identify cells. Using a CellProfiler image analysis pipeline (McQuin *et al*., 2018), a fluorescent threshold was set above background that selected mCherry-Slf1 nodes. The intensity of each node and the number of nodes per cell was output. The average number of nodes per cell was determined and plotted on a graph (Figure S6B). The average mCherry-Slf1 node intensity per cell was calculated and plotted on a graph and normalized to WT cells (Figure S6C).

### FRAP analysis

To perform FRAP experiments (Figure 3A-B & Figure 5A-B), images were captured at a single z-section on an agar pad at room temperature using a Zeiss LSM 880 laser scanning confocal microscope equipped with 100X alpha Plan-Apochromat/NA 1.46 Oil DIC M27 Elyra objective, GaAsP Detectors, and Zen Blue acquisition software. The middle focal plane of cells was chosen to bleach. After collecting 3 prebleach images, selected ROIs were bleached to <50% of the original fluorescence intensity using the laser scanner. Postbleach images were acquired for a duration long enough so that the recovery curve reached a plateau. After background subtraction and correcting for photobleaching, the data were normalized to the mean prebleach intensity of the ROI and set to 100%. The intensity just after bleaching was set to 0% so that FRAP curves represent the percent recovery of fluorescent signal. The intensity of every three consecutive postbleach time points was averaged to reduce noise. The intensity data were plotted and fitted using the exponential equation y = m1 + m2 * exp(-m3 * X), where m3 is the off-rate, using Prism 8 (GraphPad Software). The half-time of recovery was calculated using the equation t1/2 = (ln2)/m3.

### Statistical analysis

Data analysis was performed using Prism 8 (GraphPad Software). A two-tailed student’s t test was performed to determine statistical differences between two sets of data. An ANOVA was performed to determine statistical differences between sets of data.

## Abbreviations

GAP: GTPase-activating protein
PM: plasma membrane
WT: wild type
FRAP: Fluorescence recovery after photobleaching
ts: temperature-sensitive
SNARE: soluble N-ethylmaleimide-sensitive factor-attachment protein receptors

## ACKNOWLEDGEMENTS

We thank members of the Moseley laboratory for helpful discussions and comments on the manuscript; as well as the Biomolecular Targeting Core (BioMT) (P20-GM113132) and the Imaging Facility at Dartmouth for use of equipment; and Sophie Martin, Jian-Qiu Wu, Pilar Perez, and Juan Carlos Ribas for sharing yeast strains. This work was funded by grants from the National Institutes of General Medical Sciences (NIGMS) (R01GM099774 and R01GM133856) to J.B.M.

## SUPPLEMENTAL FIGURE LEGENDS

**Figure S1:**
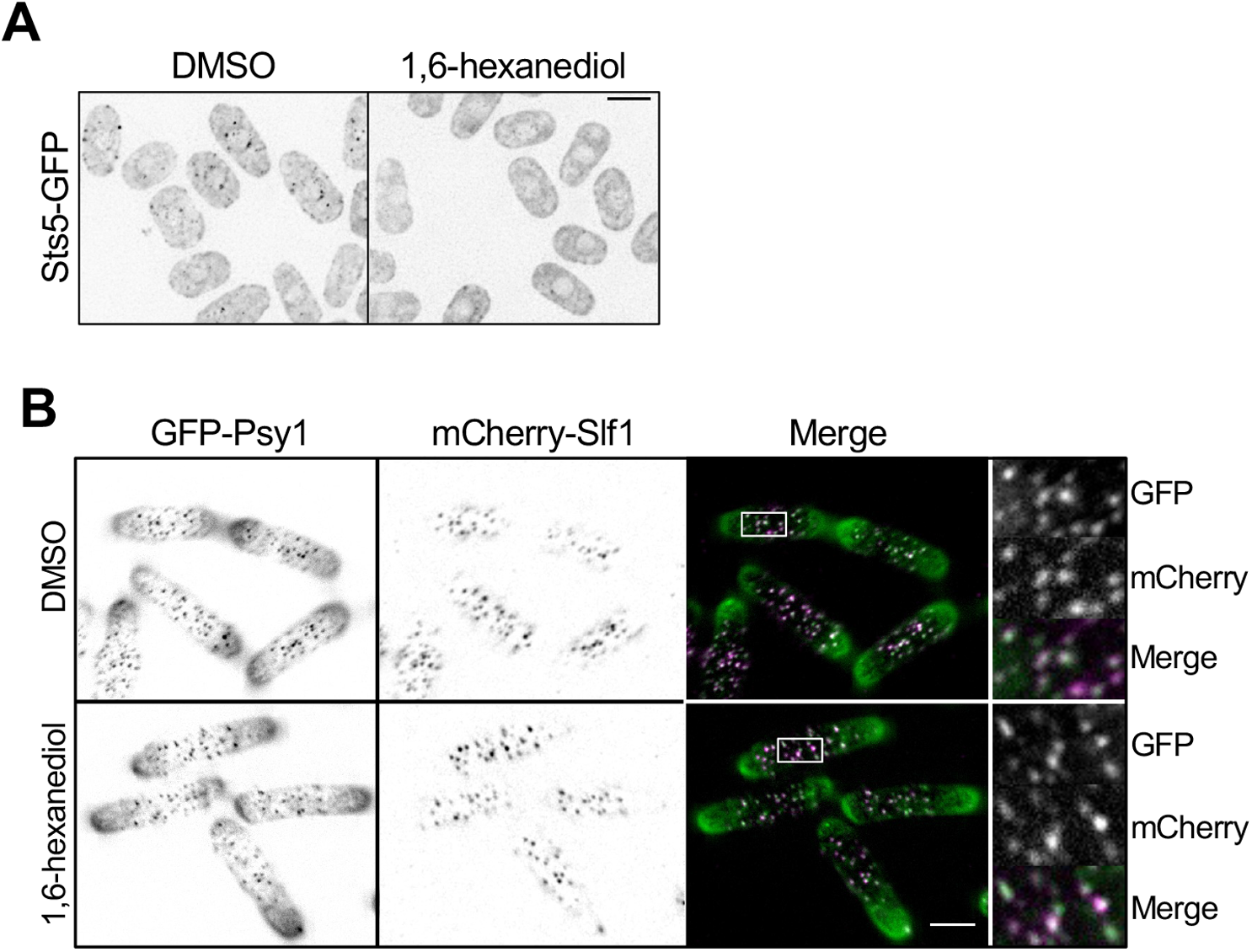
Skb1-Slf1-Psy1 nodes are stable after treatment with 1,6-hexanediol. (A) WT cells expressing P-body marker Sts5-GFP grown to saturation and then treated with 10% 1,6-hexanediol or DMSO control. (B) *GFP-psy1 mCherry-slf1* cells were treated with 10% 1,6-hexanediol or DMSO control. Images (right) are close ups of the white boxed region showing intact Skb1-Slf1-Psy1 nodes. Maximum intensity projections were created from the top half of cell z-stack images that encompass one side of the cell membrane. Bars, 4µm.

**Figure S2:**
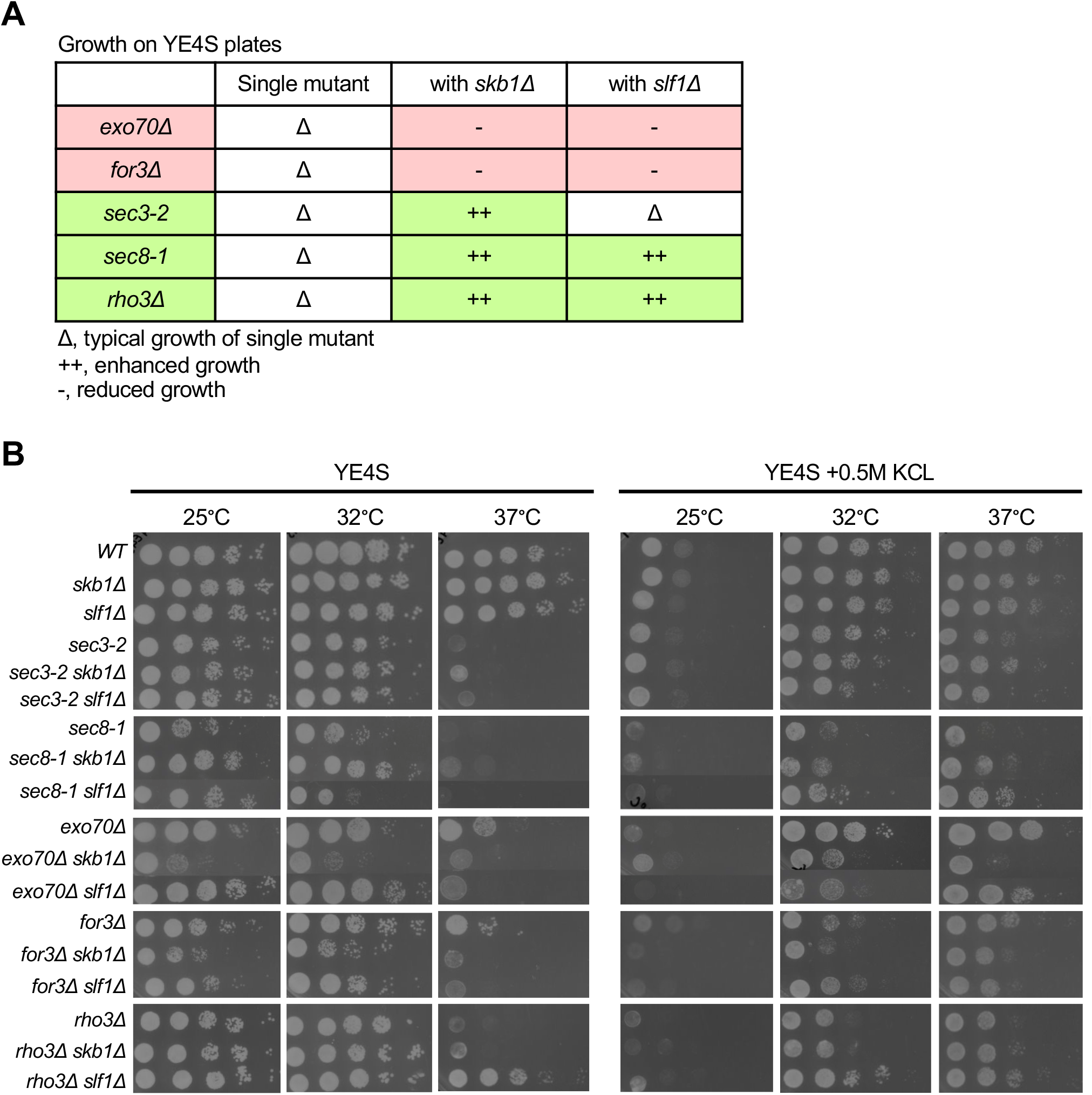
Genetic interactions between node mutants and exocytosis mutants. (A) Table summarizing growth of indicated single or double mutants shown below. Red shaded rows indicate temperature sensitive growth with double mutants compared to single mutants. Green shaded rows indicate growth suppression with double mutants. (B) The indicated strains were spotted with 10× serial dilutions onto YE4S or YE4S + 0.5M KCl plates. Plates incubated for 2-4 days prior to imaging.

**Figure S3:**
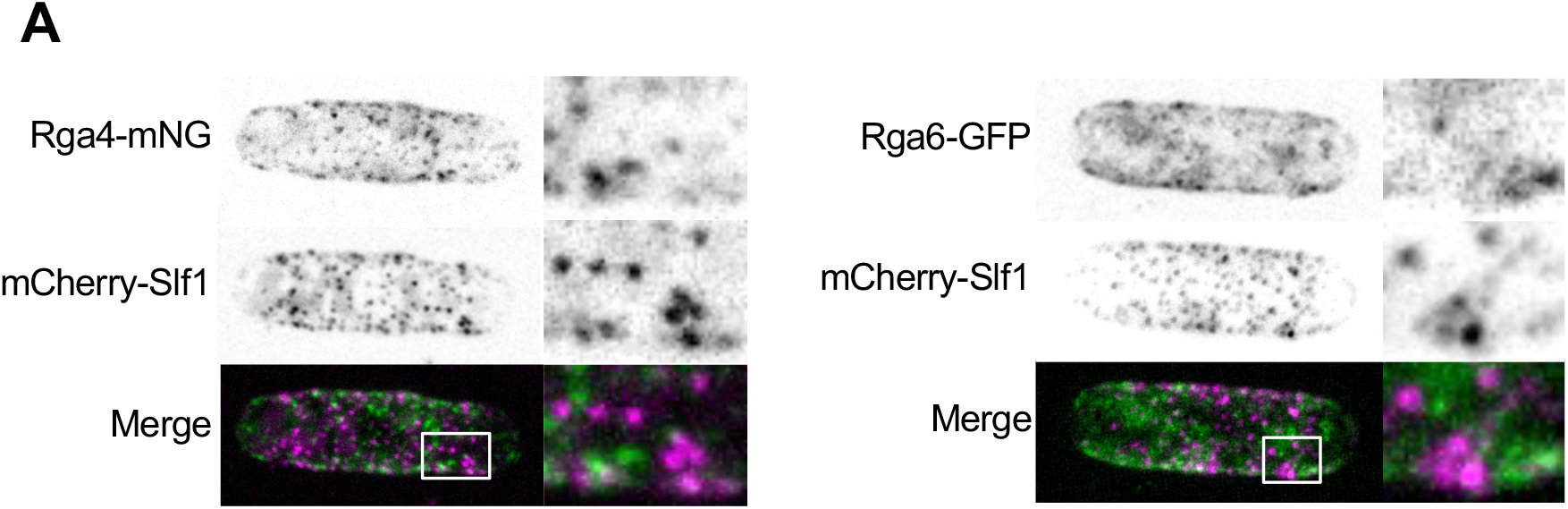
Additional analysis of Skb1-Slf1-Psy1 nodes with Rga4 and Rga6 GAPs. (A) Localization of Rga4-mNG (Left) or Rga6-GFP (Right) and mCherry-Slf1 in WT cells. Representative images with boxed region of close-up of mCherry and GFP node signals (Right panels) are shown.

**Figure S4:**
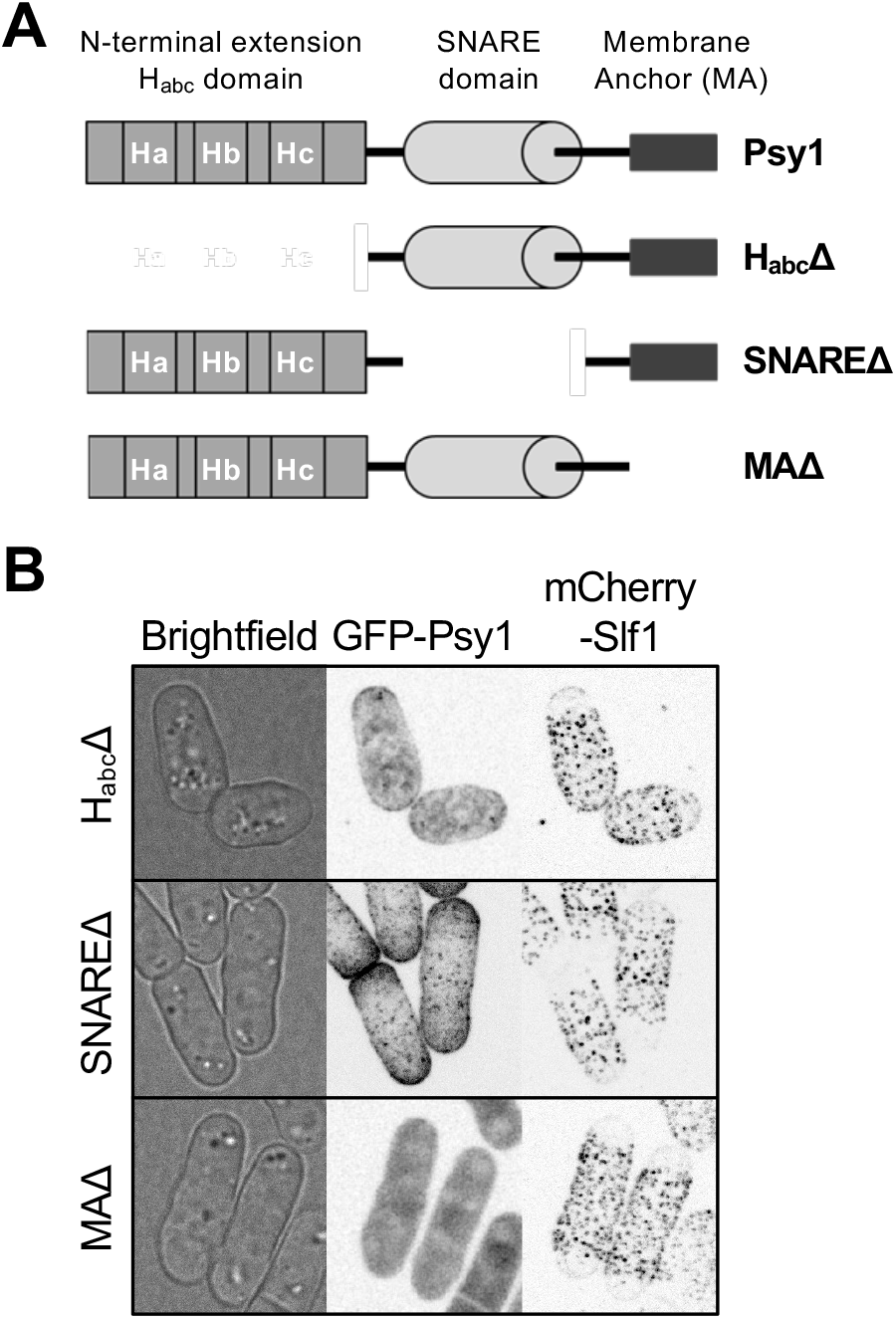
Psy1 domain deletion analysis. **(A)** Diagram of Psy1 protein domains (top). Regions included in each psy1 domain mutant is shown (below). (B) Co-localization of mCherry-Slf1 with each GFP-tagged psy1 domain mutant as listed. Images are maximum projections from z-series except for brightfield images which are single middle z-plane images.

**Figure S5:**
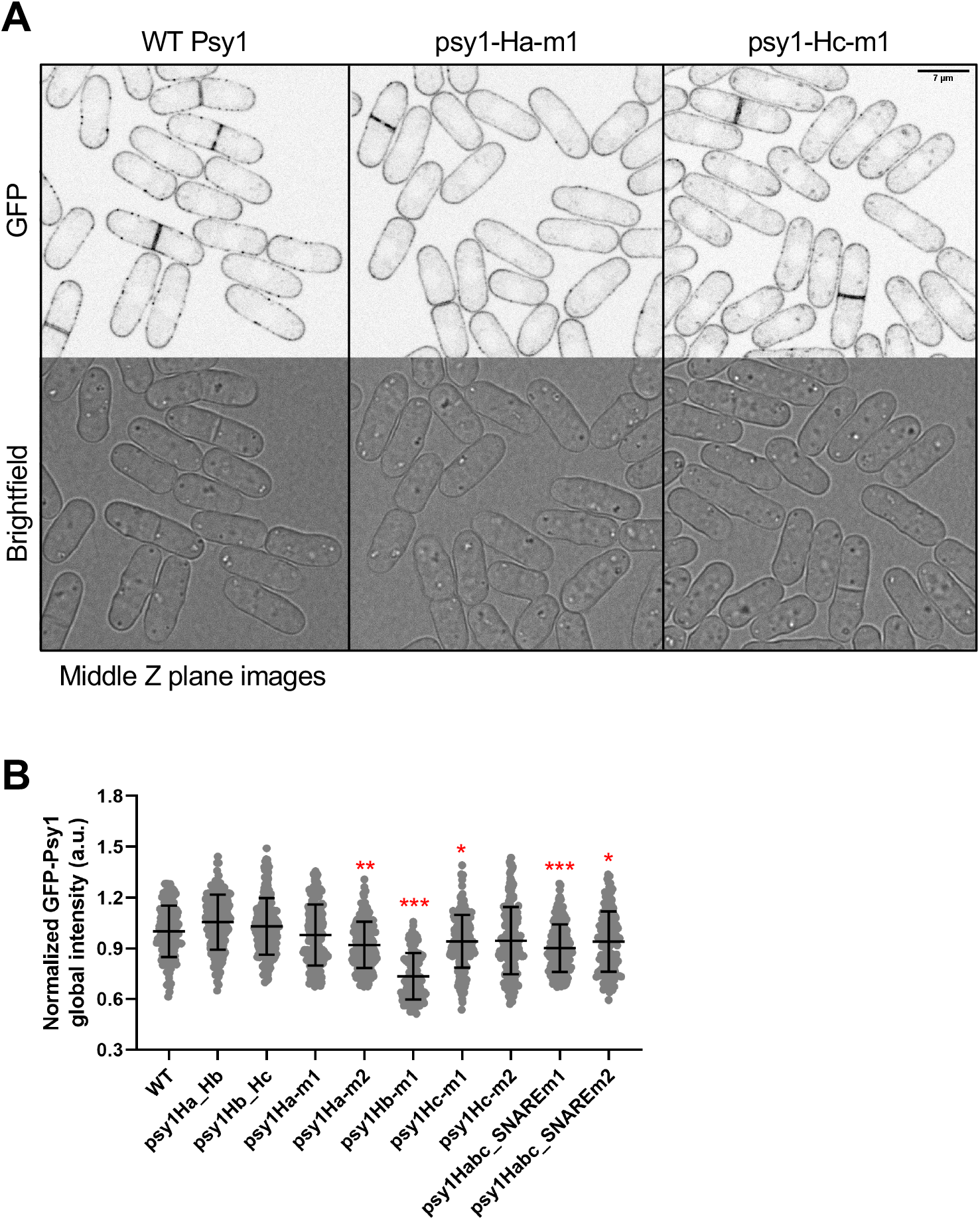
Additional characterization of GFP-tagged psy1 mutants. (A) Note the absence of nodes in GFP-psy1-Ha-m1 and GFP-psy1-Hc-m1 cells. psy1-Hc-m1 also localizes to internal puncta. Single middle z-slice images are shown. Bar, 7µm. (B) Normalized global intensity (mean ± SD) of GFP-Psy1 in indicates strains. *p < 0.05; ** p < 0.0003; and p***<0.0001 determined by ANOVA. n > 100 for each cell type.

**Figure S6:**
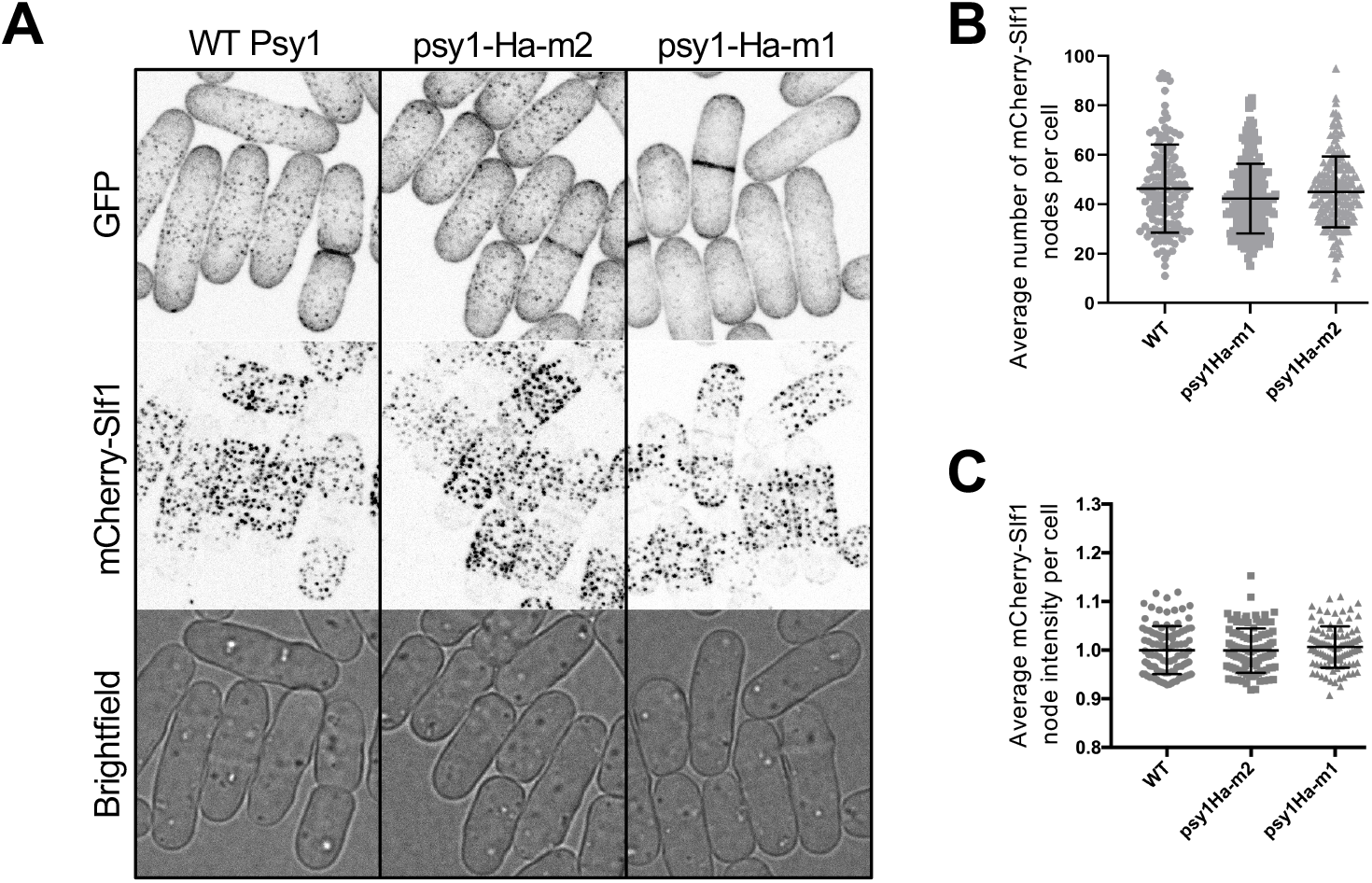
Nodes remain intact in psy1-Ha-m1 mutant. (A) Co-localization of mCherry-Slf1 with GFP-tagged Psy1, psy1-Ha-m1, or psy1-Ha-m2. Images are maximum projections from z-series except for brightfield images which are single middle z-plane images. (B) Average number of mCherry-Slf1 nodes per cell of indicated cell type (mean ± SD). No significant difference among strains determined by ANOVA. n>75 for each cell type. (C) Average mCherry-Slf1 node intensity per cell of indicated strains. mean ± SD shown. No significant difference among strains determined by ANOVA. n>75 for each cell type.

