## Supplemental Table S1 for "Sequestration of the exocytic SNARE Psy1 into multiprotein nodes reinforces polarized morphogenesis in fission yeast"

### Table S1. Yeast strains used in this study

| Strain | Genotype | Source |
|---|---|---|
| **JM6543** | *pAct1-Lifeact-mCherry::leu+ GFP-psy1::kanMX6* | This Study |
| **JM4209** | *GFP-psy1::kanMX6 mCherry-slf1::kanMX6* | This Study |
| **JM4497** | *GFP-psy1::kanMX6 mCherry-slf1::kanMX6 skb1Δ::natR* | This Study |
| **JM4207** | *GFP-Psy1::kanMX6 skb1Δ::natR* | Lab collection |
| **JM4467** | *GFP-psy1::kanMX6 slf1Δ::hphR* | This Study |
| **JM4200** | *GFP-psy1::kanMX6* | This Study |
| **JM2325** | *skb1-GFP::kanMX6 h-* | Lab collection |
| **JM1801** | *slf1-GFP::kanMX6 h+* | Lab collection |
| **JM366** | 972 *h-* | Lab collection |
| **JM906** | *skb1Δ::kanMX6 h+* | Lab collection |
| **JM909** | *skb1Δ::ura4+ ura4-D18 leu1-32* | Lab collection |
| **JM656** | *sec8-1 ura4- D18 leu1-32 h+* | Lab collection |
| **JM1449** | *exo70::kanMX6 ura4- D18 leu1-32 ade6-M21X h+* | Lab collection |
| **JM665** | *for3Δ::kanMX6 ura4-D18 leu1-32 ade6-M21X h+* | Lab collection |
| **JM6226** | *sec3-2-his5+-ura+ ade6- leu1- ura4D-18 h-* | (Bendezú *et al.,* 2012) |
| **JM6256** | *rho3Δ::kanMX6 h-* | This Study |
| **JM6057** | *skb1Δ::kanMX6 exo70Δ::kanMX6 ura4-D18* | This Study |
| **JM6262** | *skb1Δ::kanMX6 for3Δ::kanMX6 ura4-D18 h-* | This Study |
| **JM6113** | *skb1Δ::kanMX6 sec8-1 ura4-D18 leu1-32 h-* | This Study |
| **JM6313** | *rho3Δ::kanMX6 skb1Δ::natR h+* | This Study |
| **JM6374** | *sec3-2-his5+-ura+ skb1Δ::natR ura4-D18 h-* | This Study |
| **JM6112** | *slf1Δ::hphR h-* | This Study |
| **JM6127** | *slf1Δ::hphR exo70Δ::kanMX6 h-* | This Study |
| **JM6264** | *slf1Δ::hphR for3Δ::kanMX6 h-* | This Study |
| **JM6330** | *slf1Δ::hphR rho3Δ::kanMX6 h-* | This Study |
| **JM6357** | *slf1Δ::hphR sec8-1 ura4-D18 leu1-32 h+* | This Study |
| **JM6420** | *slf1Δ::kanMX6 sec3-2-his5+-ura4+ ura4-D18 leu1-32* | This Study |
| **JM6211** | *bgs4Δ::ura4+ Pbgs1+::GFP-bgs4+:leu1+ skb1Δ::NAT h+* | This Study |
| **JM4011** | *leu1-32 ura4-18 his3-1 bgs4Δ::ura4+ Pbgs1+::GFP-bgs4+:leu1+ h-* | (Cortes *et al.,* 2005) |
| **JM6210** | *bgs4Δ::ura4+ Pbgs1+::GFP-bgs4+:leu1+ slf1Δ::kanMX6 h+* | This Study |
| **JM6061** | *slf1Δ::kanMX6 sec8-mNeonGreen::hphR h+* | This Study |
| **JM6838** | *slf1Δ::hphR for3Δ::kanMX6 sec8-mNeonGreen::hphR h+* | This Study |
| **JM6802** | *slf1Δ::hphR rga4Δ::kanMX6 rga6Δ::Kan sec8-mNeonGreen::hphR h-* | This Study |

| | | |
|---|---|---|
| **JM6087** | *sec8-mNeonGreen::hphR h+* | This Study |
| **JM6398** | *skb1Δ::natR sec8-mNeonGreen::hphR* | This Study |
| **JM6641** | *skb1Δ::natR for3Δ::kanMX6 sec8-mNeonGreen::hphR h+* | This Study |
| **JM6643** | *for3Δ::kanMX6 sec8-mNeonGreen::hphR h+* | This Study |
| **JM6669** | *rga4Δ::kanMX6 rga6Δ::kanMX6 sec8-mNeonGreen::hphR h+* | This Study |
| **JM6671** | *skb1Δ::nat rga4Δ::kanMX6 rga6Δ::Kan sec8-mNeonGreen::hphR h+* | This Study |
| **JM508** | *skb1Δ::kanMX6 h-* | This Study |
| **JM2139** | *slf1Δ::hphR h+* | This Study |
| **JM6224** | *rga4Δ::kanMX6 rga6Δ::kanMX6 h-* | This Study |
| **JM6225** | *rga4Δ::kanMX6 rga6Δ::kanMX6 skb1Δ::natR* | This Study |
| **JM6465** | *slf1Δ::hphR rga4Δ::kanMX6 rga6Δ::kanMX6* | This Study |
| **JM6114** | *Pslf1-mCherry-slf1::kanMX6 rga4-mNeonGreen::hphR h-* | This Study |
| **JM6124** | *Pslf1-mCherry-slf1::kanMX6 rga6-GFP::kanMX6 h+* | This Study |
| **JM7116** | *psy1Δ::kanMX6 pDC99[Ppsy1-GFP-psy1-Tpsy1]::ura+* | This Study |
| **JM7046** | *mCherry-slf1::kanMX6 pDC99[Ppsy1-GFP-psy1ΔHabc-Tpsy1]::ura+ h+* | This Study |
| **JM7044** | *mCherry-slf1::kanMX6 pDC99[Ppsy1-GFP-psy1ΔSNARE-Tpsy1]::ura+ h-* | This Study |
| **JM7045** | *mCherry-slf1::kanMX6 pDC99[Ppsy1-GFP-psy1ΔMA-Tpsy1]::ura+ h-* | This Study |
| **JM7276** | *psy1Δ::kanMX6 pDC99[Ppsy1-GFP-psy1(51QEID54→51GGGG54)-Tpsy1 ]::ura+* | This Study |
| **JM7359** | *psy1Δ::kanMX6 pDC99[Ppsy1-GFP-psy1(91MQLPPD96→91GGGGSG96)-Tpsy1]::Ura+* | This Study |
| **JM7328** | *psy1Δ::kanMX6 pDC99[Ppsy1-GFP-psy1(E20A, E23A, D26A, H27A, D30A)-Tpsy1]::Ura+ h-* | This Study |
| **JM7329** | *psy1Δ::kanMX6 pDC99[Ppsy1-GFP-psy1(R33A, E36A, D37A, R41A, M44A)-Tpsy1]::Ura+* | This Study |
| **JM7360** | *psy1Δ::kanMX6 pDC99[Ppsy1-GFP-psy1(R145A, E148A, D149A, D150A, F151A)-Tpsy1]::Ura+* | This Study |
| **JM7421** | *psy1Δ::kanMX6 pDC99[Ppsy1-GFP-psy1(V163A, L168A, R173A)-Tpsy1]::Ura+* | This Study |
| **JM7422** | *psy1Δ::Kan pDC99[Ppsy1-GFP-psy1(D98A, K103A, K111A, K112A, D115A)-Tpsy1]::Ura+ h-* | This Study |
| **JM7423** | *psy1Δ::kanMX6 pDC99[Ppsy1-GFP-psy1(R63A, H64A, E66A, Y68A, D71A)-Tpsy1]::Ura+* | This Study |
| **JM7424** | *psy1Δ::kanMX6 pDC99[Ppsy1-GFP-psy1(R118A, H119A, L121A, K125A, R128A, R136A, R137A)-Tpsy1]::Ura+ h-* | This Study |
| **JM6888** | *sec8-tdTomato-kanMX6 h+* | This Study |
| **JM7430** | *sec8-tdTomato-kanMX6 psy1Δ::kanMX6 pDC99[Ppsy1-GFP-psy1(E20A, E23A, D26A, H27A, D30A)-Tpsy1]::Ura+* | This Study |
| **JM7445** | *sec8-tdTomato-kanMX6 rga4Δ::kanMX6 rga6Δ::kanMX6 psy1Δ::kanMX6 pDC99[Ppsy1-GFP-psy1(E20A, E23A, D26A, H27A, D30A)-Tpsy1]::Ura+* | This Study |
| **JM7446** | *sec8-tdTomato-kanMX6 rga4Δ::kanMX6 rga6Δ::kanMX6* | This Study |
| **JM6025** | *sts5-mNeonGreen::HphR ura4D-18 leu1-32* | This Study |
| **JM7185** | *psy1Δ::kanMX6 pDC99[Ppsy1-GFP-psy1-Tpsy1]::Ura+ Pslf1-mCherry-slf1::kanMX6* | This Study |

| | | |
|---|---|---|
| **JM7372** | *psy1Δ::kanMX6 pDC99[Ppsy1-GFP-psy1(E20A, E23A, D26A, H27A, D30A)-Tpsy1]::Ura+ Pslf1-mCherry-slf1::kanMX6* | This Study |
| **JM7373** | *psy1Δ::kanMX6 pDC99[Ppsy1-GFP-psy1(R33A, E36A, D37A, R41A, M44A)-Tpsy1]::Ura+ Pslf1-mCherry-slf1::kanMX6* | This Study |
