## Supplemental Table S2 for "Sequestration of the exocytic SNARE Psy1 into multiprotein nodes reinforces polarized morphogenesis in fission yeast"

### Table S2. Plasmids used in this study

| Strain | Genotype | Source |
|---|---|---|
| **pJM210** | pRep3x | Lab Collection |
| **pJM482** | pREP3x-6His-skb1 | Deng *et al.,* 2013 |
| **pJM1551** | pDC99-Ppsy1-GFP-psy1-Tpsy1 | This Study |
| **pJM1567** | pDC99-Ppsy1-GFP-psy1Δ$H_{abc}$(Δ11-170aa)-Tpsy1 | This Study |
| **pJM1568** | pDC99-Ppsy1-GFP-psy1ΔSNARE(Δ180-250aa)-Tpsy1 | This Study |
| **pJM1569** | pDC99-Ppsy1-GFP-psy1ΔMA(Δ252-283aa)-Tpsy1 | This Study |
| **pJM1591** | pDC99-Ppsy1-GFP-psy1(51QEID54 →51GGGG54)-Tpsy1<br>Ha_Hb mutant | This Study |
| **pJM1599** | pDC99-Ppsy1-GFP-psy1(91MQLPPD96 → 91GGGGSG96)-Tpsy1<br>Hb_Hc mutant | This Study |
| **pJM1597** | pDC99-Ppsy1-GFP-psy1(E20A, E23A, D26A, H27A, D30A)-Tpsy1<br>Ha-m1 mutant | This Study |
| **pJM1598** | pDC99-Ppsy1-GFP-psy1(R33A, E36A, D37A, R41A, M44A)-Tpsy1<br>Ha-m2 mutant | This Study |
| **pJM1600** | pDC99-Ppsy1-GFP-psy1(R145A, E148A, D149A, D150A, F151A)-Tpsy1<br>Habc_SNARE-m1 mutant | This Study |
| **pJM1601** | pDC99-Ppsy1-GFP-psy1(V163A, L168A, R173A)-Tpsy1<br>Habc_SNARE-m2 mutant | This Study |
| **pJM1604** | pDC99-Ppsy1-GFP-psy1(R63A, H64A, E66A, Y68A, D71A)-Tpsy1<br>Hb-m1 mutant | This Study |
| **pJM1603** | pDC99-Ppsy1-GFP-psy1(D98A, K103A, K111A, K112A, D115A)-Tpsy1<br>Hc-m1 mutant | This Study |
| **pJM1605** | pDC99-Ppsy1-GFP-psy1(R118A, H119A, L121A, K125A, R128A, R136A, R137A)-Tpsy1<br>Hc-m2 mutant | This Study |
